# Phylogenomics within the Anthonotha clade (Detarioideae, Leguminosae) reveals a high diversity in floral trait shifts and a general trend towards organ number reduction

**DOI:** 10.1101/511949

**Authors:** Dario I. Ojeda, Erik Koenen, Sandra Cervantes, Manuel de la Estrella, Eulalia Banguera-Hinestroza, Steven B. Janssens, Jeremy Migliore, Boris Demenou, Anne Bruneau, Félix Forest, Olivier J. Hardy

**Affiliations:** Department of Ecology and Genetics, University of Oulu, PO Box 3000, Oulu FIN-90014, Finland; Norwegian Institute of Bioeconomy Research, 1430 Ås, Norway; Institute of Systematic Botany, University of Zurich, Zürich, Switzerland; Comparative Plant and Fungal Biology Department, Royal Botanical Gardens, Kew, Richmond, TW9 3DS, UK; Departamento de Botánica, Ecología y Fisiología Vegetal, Facultad de Ciencias, Campus de Rabanales, Universidad de Córdoba, 14071, Córdoba, Spain; Evolutionary Biology and Ecology Unit, CP 160/12, Faculté des Sciences, Université Libre de Bruxelles, Av. F. D. Roosevelt 50, B-1050 Brussels, Belgium; Botanic Garden Meise, Nieuwelaan 38, BE-1860 Meise, Belgium; Institut de recherche en biologie végétale and Département de Sciences Biologiques, Université de Montréal, 4101 Sherbrooke est, Montréal, QC H1X 2B2, Canada

**Keywords:** Berlinia clade, flower evolution, papillose conical cells, petal number, petal identity, phylogenomics, target enrichment

## Abstract

Detarioideae is well known for its high diversity of floral traits, including flower symmetry, number of organs, and petal size and morphology. This diversity has been characterized and studied at higher taxonomic levels, but limited analyses have been performed among closely related genera with contrasting floral traits due to the lack of fully resolved phylogenetic relationships. Here, we used four representative transcriptomes to develop an exome capture bait for the entire subfamily and applied it to the Anthonotha clade using a complete data set (61 specimens) representing all extant floral diversity. Our phylogenetic analyses recovered congruent topologies using ML and Bayesian methods. The genus *Anthonotha* was recovered as monophyletic contrary to the remaining three genera *(Englerodendron, Isomacrolobium* and *Pseudomacrolobium),* which form a monophyletic group sister to *Anthonotha.* We inferred a total of 35 transitions for the seven floral traits (pertaining to flower symmetry, petals, stamens and staminodes) that we analyzed, suggesting that at least 30% of the species in this group display transitions from the ancestral condition reconstructed for the Anthonotha clade. The main transitions were towards a reduction in the number of organs (petals, stamens and staminodes). Despite the high number of transitions, our analyses indicate that the seven characters are evolving independently in these lineages. Petal morphology is the most labile floral trait with a total of seven independent transitions in number and seven independent transitions to modification in petal types. The diverse petal morphology along the dorsoventral axis of symmetry within the flower is not associated with differences at the micromorphology of petal surface, suggesting that in this group all petals within the flower might possess the same petal identity at the molecular level. Our results provide a solid evolutionary framework for further detailed analyses of the molecular basis of petal identity.

## 1. Introduction

### 1.1. Flower diversity in the Detarioideae and the Anthonotha clade

Legumes are well known for their diversity in flower morphology. The group has diversified into a diverse array of flower arrangements, flower symmetry and number of organs within each of the whorls (Tucker, 2003). The best known flower arrangement in legumes (the papilionoid or “pea-like” flower) with distinctive petal types arranged along the dorsoventral axis of the flower is characteristic of most taxa within subfamily Papilionoideae. This arrangement consists of an adaxial petal (also referred to as dorsal, banner or standard petal), two lateral petals (wings), and two abaxial petals (also known as ventral or keel) (LPWG, 2017). This flower arrangement is considered highly specialized and its coevolution with bees is suggested as one of the drivers of diversification in this group (Cronk and Moller, 1997). Most of our current knowledge on flower evolution and its molecular basis has been centered in this subfamily. The remaining five subfamilies have also evolved diverse arrays of flower diversity in relation to a diverse range of pollinators (Banks and Rudall, 2016; Bruneau et al., 2014; Tucker, 2003; Zimmerman et al., 2017).

The pantropical subfamily Detarioideae comprises 81 genera and ca. 760 species, with its highest diversity in tropical Africa and Madagascar (58% of species) (de la Estrella et al., 2018, 2017). The group contains large trees of ecological importance in tropical environments (e.g. the Miombo forest in east Africa) (Ryan et al., 2016), but also several species of economic importance as food source, timber, oils, resins, as well as ornamentals (Langenheim, 2003). Detarioideae is well known for its high level of flower diversity, regarding for example the symmetry, floral arrangement and size, and number of organs per whorl (LPWG, 2017). The entire range of floral diversity encountered in present-day legumes, with the notable exception of the specialized papilionoid flower, is encompassed in this subfamily (Bruneau et al., 2014). This diversity is also visible at the flower developmental (ontogeny) level, where the final number and arrangement of floral organs at anthesis is achieved via alternative mechanisms (Bruneau et al., 2014; Tucker, 2002a, 2000). The plasticity of some of these floral traits has been documented at multiple levels, among major lineages, between closely related genera (Bruneau et al., 2014) and within species (Tucker, 2002a, 2002b).

One of the most extreme cases of variation of floral traits within Detarioideae (e.g. petal and stamen numbers) is reported in the Berlinia clade, where plasticity of these traits is observed within the same species and within flowers of the same individual (Breteler, 2011, 2010, 2008). In addition to the diversity in organ number (merism), the Berlinia clade also displays variation in petal size and arrangements within the dorsoventral axis of the flower. Most genera display a large adaxial (dorsal) petal in the flower (e.g. *Gilbertiodendron* and *Berlinia* Fig. 1A and D) with additional lateral and abaxial (ventral) petals of different size and shape (Bruneau et al., 2014; de la Estrella and Devesa, 2014a; LPWG, 2017; Mackinder and Pennington, 2011). Within the Berlinia clade, the Anthonotha clade, which comprises a group of three closely related genera with contrasting flower symmetry, is particularly diverse in its flower morphology. Some species of *Anthonotha* (Fig. 1G) and *Isomacrolobium* (Fig. 1F) display modifications in petal arrangement, whereby the dorsal, lateral and abaxial petals all possess a different morphology, or where the adaxial and lateral petals possess the same morphology while the abaxial one has a unique morphology. In contrast, in *Englerodendron* (Fig. 1E), a genus of four species with actinomorphic (radial symmetry) flowers (Breteler, 2006), all petals have the same morphology, and they resemble either the adaxial or the lateral petals of the zygomorphic and closely related *Anthonotha* and *Isomacrolobium.* These differences in petal size and morphology along the dorsoventral axis of the flower might suggest distinct petal identities at the molecular level, but this remains to be tested.

**Fig. 1.**
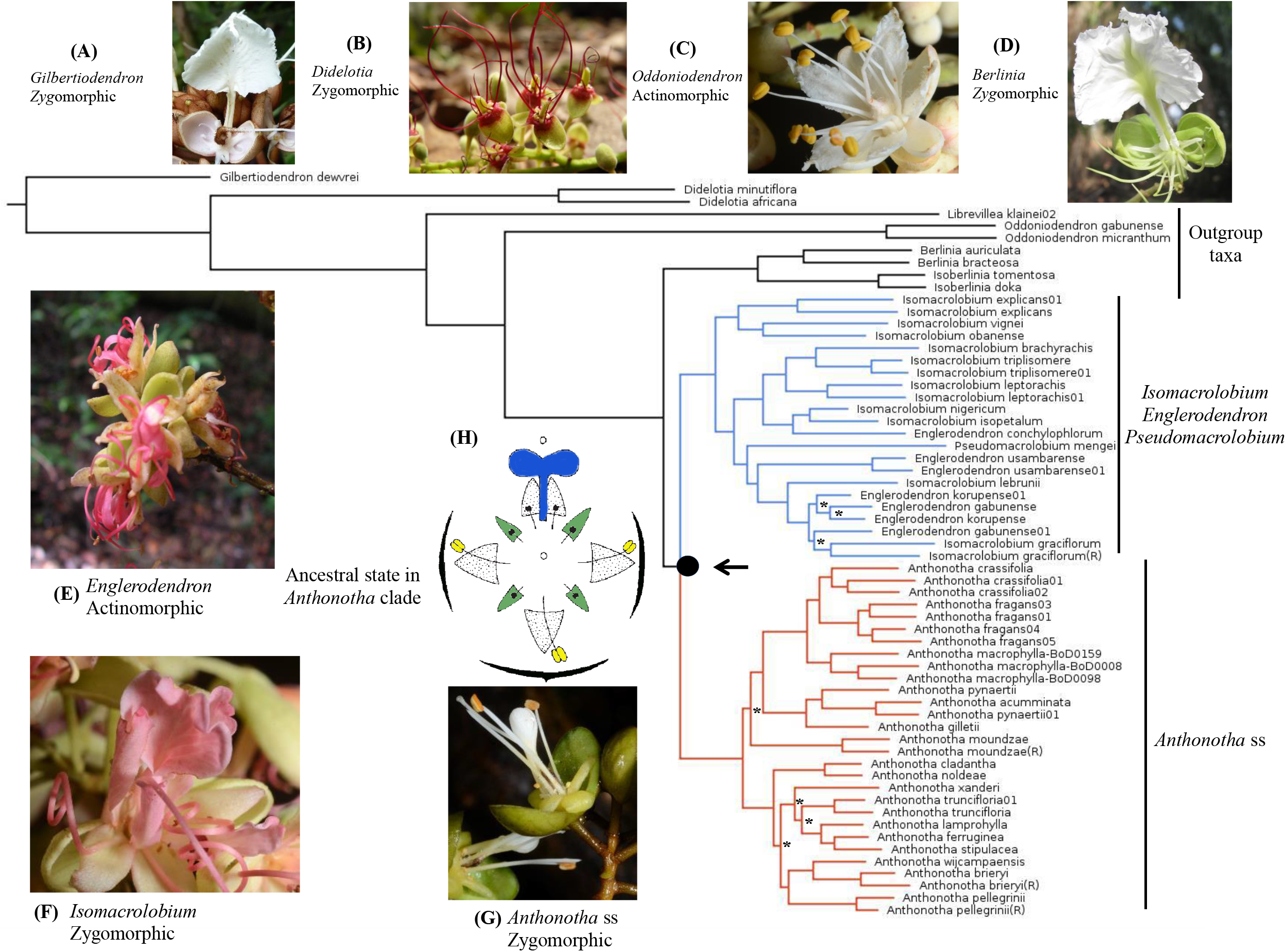
Phylogenetic reconstruction of the Berlinia clade obtained from individual and concatenated analyses using ML and Bayesian inference. The arrow indicates the location of the ancestral state reconstruction within the focal group *(Anthonotha* clade). Flower diversity in the genera sampled in the Berlinia clade: (A) *Gilbertiodendron obliquum* (Jongkind 9972), (B) *Didelotia africana* (XvdB_3), (C) *Oddoniodendron micranthum* (Bidault_MBG), (D) *Berlinia grandiflora* (FOLI092H), (E) *Englerodendron korupense* (XvdB_741 −8), (F) *Isomacrolobium* aff. *triplisomere* (Bidault_2214_EB_3234), (G) *Anthonotha macrophylla* (Bidault_1246_EB_9850), and (H) schematic representation of the flower in the amcestral state (A. *macrophylla).* Petal types (identity based on their position within the dorsoventral axis within the flower) are in color blue (adaxial petal) and in green (lateral and abaxial petals). Stamens are indicated in yellow and staminodes in black. * = Nodes with < 85 bootstrap support obtained from the concatenated and individual RAxML analyses. Drawing modified from fig. 15 in Breteler (2010: 84). Copyright Meise Botanic Garden and Royal Botanical Society of Belgium. Photo credits: (A) Carel Jongkind, (C, F, G) Ehoarn Bidault, (D) Olivier Hardy, (B, E) Xander van der Burgt.

### 1.2 The molecular basis of petal identity and its association with petal micromorphology

The molecular basis of the of petal identity has been well characterized in papilionoids, where specific transcription factors of *CYCLOIDEA* have been identified in *Lotus* (Feng et al., 2006; Wang et al., 2010; Xu et al., 2013) and *Pisum* (Wang et al., 2008). In these two genera, the differences in petal shape along the dorsoventral axis and petal symmetry reflect differences in gene expression domains of three copies of *CYCLOIDEA.* This results in three domains of identity: adaxial (dorsal), lateral and abaxial (ventral) identities. These domains of identity within the flower are also reflected at the micromorphological level on each petal, where specific epidermal types are associated with each petal identity (Feng et al., 2006; Ojeda et al., 2012, 2009). For instance, the dorsal petal identity is conferred by the activity of two copies of *CYCLOIDEA (LjCYC2*and *LjCYC1*) in *Lotusjaponicus* L. and the petal surface is characterized by papillose conical cells (PCS) (Feng et al., 2006; Xu et al., 2013). However, these molecular analyses have been concentrated in the Papilionoideae and there is a lack of understanding of the molecular basis of petal identity in the other subfamiles of the Leguminosae. The characterization of petal micromorphology and its relation to petal identity also has been focused mainly on the Papilionoideae. Only two Detarioideae genera *(Brownea, Tamarindus)* have been analyzed to date (Ojeda et al., 2009) and neither showed micromorphological differentiation among petal types. However, it is unclear whether this lack of micromorphological differentiation applies more generally to other members of this subfamily. Other subfamilies are poorly represented in our understanding of floral genetics in legumes and with this study we hope to bring awareness of the potential use of the Detarioideae as a “model clade” (Chanderbali et al., 2016) to further improve our understanding of flower evolution in Leguminosae.

### 1.3 Phylogenetic relationships of Anthonotha and closely related genera in the Berlinia clade

The Berlinia clade (tribe Amherstieae) (de la Estrella et al., 2018) as circumscribed by Bruneau et al (2008) comprises 17 genera and some ca. 180 species that occur exclusively in Africa. Previous studies based on morphology and a few molecular markers have identified major lineages within this group, e.g. the “babjit” group (Wieringa, 1999), which also has been recovered in more recent phylogenetic analyses (de la Estrella et al., 2018, 2014). However, generic relationships have not been fully resolved and a limited number of species have been included for most genera within the Berlinia clade in previous studies (Bruneau et al., 2008; de la Estrella et al., 2014; Mackinder et al., 2013). Within the Berlinia clade, the genus *Anthonotha* and its two close relatives, *Isomacrolobium* and *Englerodendron,* is potentially the most problematic group that remains to be studied because several studies have suggested *Anthonotha* might not be monophyletic (Breteler, 2008). The most recent taxonomic analyses of the *Anthonotha* clade have been entirely based on morphological data, and it is not clear whether the morphological traits used to circumscribe these genera reflect monophyletic lineages (Breteler, 2011, 2010, 2008, 2006; van der Burgt et al., 2007). Despite the remarkable flower diversity among the 17 genera of the Berlinia clade, and in particular within the *Anthonotha* clade, the lack of phylogenetic resolution at the generic level found in previous studies has hampered in-depth analyses of floral evolution. In order to better resolve relationships among closely related genera within Detarioideae, as well as within genus, we developed a target enrichment (exome capture) for the entire subfamily by selecting orthologues shared among four representative transcriptomes. We combined the design of this capture bait set with a complete sampling of *Anthonotha* and its closely related genera, representing a group of ca. 35 species with high diversity in floral traits.

### 1.4 Objectives of this study

The aims of this study were to: 1) reconstruct the relationships of *Anthonotha* and its most closely related genera *(Isomacrolobium* and *Englerodendron)* within the Berlinia clade using a complete sampling representing all extant floral diversity, 2) determine the ancestral states of seven of the most labile floral traits within the Anthonotha clade and identify major transitions from the ancestral floral state reconstructed, 3) test for correlations in the evolution of the floral traits investigated in this group, and 4) establish whether changes in petal morphology and position within the dorsoventral axis of the flower are associated with differences in petal identity, as measured by differences in petal surface micromorphology.

## 2. Materials and methods

### 2.1 Selection of species and transcriptome sequencing

For the transcriptome analysis, we selected four species *(Anthonotha fragans* (Baker f.) Exell & Hillc., *Afzelia bella* Harms, *Copaifera officinalis* (Jacq.) *L*., and *Prioria balsamifera* (Vermoesen) Breteler) representing the major clades within Detarioideae (de la Estrella et al., 2018). Three were generated in this study and the fourth was obtained from a previous study on *Copaifera officinalis* Jacq. (Matasci et al., 2014). For each species we collected young leaves from seedlings growing under nursery conditions. RNA was extracted using the Pure Link™ Plant RNA Reagent (Invitrogen) following the manufacturer’s protocol with DNAse I Amp grade (Invitrogen). RNA quantity and quality were checked using a Qubit^R^ 2.0 Invitrogen (Life Technologies) and with an agarose gel at 2%, respectively. Each sample was later enriched for mRNA using the NEXTflex™ Poly(A) bead capture (BioScientific). RNA libraries were prepared for each species with the NEXTflex™ Rapid Directional RNA-seq kit following the manufacturer’s protocol and ~25 million reads (150 bp paired end sequences) were generated per library on a NextSeq Illumina sequencer at GIGA (Grappe Interdisciplinaire de Génoprotéomique Appliquée) at the Université de Liège. Raw reads were first analyzed with FastQC (http://www.bioinformatics.bbsrc.ac.uk/projects/fastqc) and later cleaned with Trimmomatic v. 036 (Bolger et al., 2014) with settings ILLUMINACLIP:TruSeq3-PE.fa:20:30:10 LEADING:3 TRAILING:3 SLIDINGWINDOW:4:15 MINLEN:36. Trinity v. 2.2 (Grabherr et al., 2011) was used for *de novo* assembly using the default parameters. Raw reads for each species were assembled separately. Transcriptome assembly statistics and quality were assessed with rnaQUAST v. 1.4 (Bushmanova et al., 2016). To assess completeness of the transcriptomes, the Benchmarking Universal Single-Copy Orthologs (BUSCO ver. 2) was run on each species separately using the embryophyta odb9 database (Simão et al., 2015). Statistics of the four assemblies are described on Table S1. Fastq sequences from the three transcriptomes are deposited in the NCBI Bioproject PRJNA472454 (SUB4060777, SUB4060776, SUB4060712).

### 2.2. Target loci selection and bait design

For each species we identified open reading frames (ORFs) in the assembled transcriptome contigs using TransDecoder v2.1.0 (Haas et al., 2013) with defaults parameters. Coding regions with at least 100 amino acids were first predicted and extracted. An additional retention criteria search was used to retain ORFs with homology blasts on legumes using a BlastP search against a database containing the proteins from the complete genomes of six legume species *(Cajanus cajan* (L.) Millsp., *Cicer arietinum L., Glycine max* (L.) Merr., *Medicago truncatula* Gaertn., *Phaseolus vulgaris* Wall. and *Vigna radiata* (L.) R. Wilczek) (Data S1). We identified putatively single copy (or low copy number) genes with a CD-HIT (Li and Godzik, 2006) to cluster highly similar proteins within each species, followed by a SelfBLAST step to discard proteins with multiple hits. We later identified orthologues among the four transcriptomes using reciprocal best blast hits using a BlastP with a cut-off value of 1 -evalue 1e-10. We found from 32,129 to 46,510 ORFs with BlastP hits with the Pfam and the legume genomes (Table S1).

Selected orthologues were first aligned with MAFFT v.7 (Katoh and Kuma, 2002) and trimmed with BMGE (Criscuolo and Gribaldo, 2010). Individual phylogenetic trees were reconstructed from each identified shared orthologue using RAxML v. 8.2.9 (Stamatakis, 2014) and only genes with a congruent topology to the known phylogenetic relationships among the four species were retained (de la Estrella et al., 2018, 2017). We generated two data sets, one with the recovered orthologues considering the four transcriptomes and the tree topology, and another set excluding *Afzelia bella* (with lowest number of contigs in the assembled transcriptomes). Orthologs with >80 *%* homology and at least 300 bp in length were selected from these two data sets. Putative intron/exon boundaries within these selected genes were predicted using the *A. fragans* assembled contigs as a reference and a database of three legume genomes *(M. truncatula, G. max* and *C. aurietinum)* using a custom-made python script. Only exons >100 bp were retained along with the 5’ and 3’ UTR sequences attached to the exons of the flanking regions. Each exon was treated separately for bait design to prevent location of the baits in the intron/exon boundaries. We selected 289 target loci (genes), representing 1,058 contigs with a total of 1,021 putative exons and a total size of 359,269 bp. The selected sequences were subjected to a RepeatMasker (www.repeatmasker.org) analysis (Smit et al., 2015) using the Fabaceae repeats database from the Michigan State University Plant Repeat Databases (http://plantrepeats.plantbiology.msu.edu/downloads.html) with default settings. The probes were further *in silico* screened separately against both the *G. max* and *M. truncatula* genomes. Probes with soft-masking sequences and with non-unique hits in the two reference genomes were excluded from the final bait design. The selected target regions were used to select orthologues and to develop a target enrichment bait suitable for the entire subfamily. The final set of genes selected were used to design an exome capture bait (Arbor Biosciences, MI, USA) with a 3x tiling of 120 bp RNA baits generating a total of 6,565 probes. This probe was applied to the Anthonotha clade, the focal group of our study, together with selected genera of the Berlinia clade.

### 2.3 Taxon sampling

The focal group of the study (Anthonotha clade) consists of c. 35 species currently classified based in morphology into four genera: *Anthonotha* (17 species), *Englerodendron* (4 species), *Isomacrolobium* (12 species) and the monospecific *Pseudomacrolobium (P. mengei* Hauman). Our sampling included 61 samples representing all species within the Anthonotha clade (except *Isomacrolobium sargosii* (Pellegr.) Aubrév. & Pellegr. and *I. hallei* Aubrév.) (Table S2). This sampling included several individuals for widely distributed species and replicates for four species. In order to represent the closest genera of our focal group, we also included eight of the 17 genera within the Berlinia clade. These eight genera have been previously identified as the closest related lineages of our focal group (de la Estrella et al., 2018, 2017, 2014).

### 2.4 DNA extraction, library preparation and hybridization

Total genomic DNA was extracted from leaf tissue (25-35 mg) from herbarium specimens or silica gel samples (Table S2) using the CTAB modified protocol (Doyle and Doyle, 1987) and the QIAquick PCR Purification Kit (Qiagen, Venlo, Netherlands), followed by Qubit^®^ 2.0 Fluorometer quantification (Life Technologies, Invitrogen, Foster City, USA). We used a modification of the cost-effective protocol for plastome capture for library preparation (Mariac et al., 2014). Briefly, DNA extracts were sheared using a Bioruptor^®^ Pico (Diagenode SA., Liège, Belgium) to yield sonicated fragments of around ca. 400 bp. Sheared and sized DNA was then repaired and tagged using 6-bp barcodes for multiplexing all the samples (Rohland and Reich, 2012) after AMPure XP bead-based sample clean-up steps (Agencourt Bioscience, Beckman Coulter, Brea, USA). Hybrid enrichment was performed on pools of 48 samples per reaction following the MYbaits v 3.0.1 protocol, with 24 h of hybridization, a high stringency post-hybridization wash and a final amplification involving 17 PCR cycles on a StepOnePlus (Applied Biosystems, Foster City, USA). Paired-end sequencing (2 × 150 bp) was performed on an Illumina NextSeq with reagent kit V2 at the GIGA platform (Liège, Belgium), assigning 400,000 million reads/sample.

### 2.5 Assembly of captured sequences and recovery of orthologues

Raw reads were first analyzed with FastQC and later cleaned with Trimmomatic v. 036 (Bolger et al., 2014) using the same conditions as described above. Reads were demultiplexed using the software Sabre (https://github.com/najoshi/sabre) and mapped to the bait reference using BWA with the bwa-mem algorithm (Li and Durbin, 2010). We then used Samtools (Li et al., 2009) and bedtools (Quinlan and Hall, 2010) on the mapped reads to determine capture success and coverage for each sample. Reads for each sample were later assembled *de novo* using SPAdes ver. 3.9 (Bankevich et al., 2012). We chose the genomic version of SPAdes (instead of rnaSPAdes), which reduces the number of isoforms on the assemblies. This assembler uses iterative k-mer lengths during the assembly allowing the reconstruction of contigs from the long reads (150 bp) we employed in this study. The contigs generated represent a consensus sequence of each captured region, thus indels and heterozygotes are not included, as the consensus sequence represents the most common allele. We used the cds and peptides of *A. fragans* reference (Data S2) and a custom-made Python script to select the corresponding contigs with significant hits (e-value 1e^-10^) using Blast (Altschul et al., 1997) and Blat (Kent, 2002). We further identified orthologues on the selected contigs using the strategy developed by Yang and Smith (2014) (https://bitbucket.org/yangya/phylogenomic_dataset_construction).

Briefly, we reduced redundancy of cluster using CD-HIT (Li and Godzik, 2006) (99%, threshold, word size =10). After that we performed an all-by-all blast on all the samples and later filtered with a hit fraction cut-off of 0.5. We then used MCL (Van Dongen, 2000) with an inflation value of 1.4 to reduce identified clusters in the samples. Clusters with less than 1000 sequences were aligned with mafft (--genafpair --maxiterate 1000) (Katoh and Kuma, 2002), 0.1 minimal column occupancy and tree inference was generated with RAxML v. 8.2.9 (Stamatakis, 2014). We used PASTA (Mirarab et al., 2015) for larger clusters, minimal column occupancy of 0.01 and trees were inferred using fasttree (Price et al., 2009). Paralog sequences were pruned using the strict 1to1strategy (Yang and Smith, 2014), which searches for homolog sequences that are strictly one-to-one correspondence among samples. The concatenated alignment was visually inspected and formatted with AliView (Larsson, 2014) and summary statistics obtained using AMAS (Borowiec, 2016). Raw sequence reads are deposited in the NCBI SRA XXXXXX.

### 2.6 Phylogenomic analysis using gene tree (individual orthologues) and supermatrix (concatenation) approaches

Phylogenetic analyses were performed using maximum likelihood approach (ML) as implemented in RAxML v. 8.2.9 (Stamatakis, 2014) on each separate orthologue alignment using the GTRCAT model with -f a flags, 1000 bootstrap replicates and default settings. Additional analyses were performed using a supermatrix (concatenated alignment) by ML as described above and with a Bayesian approach using MrBayes 3.2.6 (Huelsenbeck and Ronquist, 2001; Ronquist and Huelsenbeck, 2003) as implemented in CIPRESS Gateway (Miller et al., 2010) using four chains, two runs of 5,000,000 generations with the invgamma rate of variation and a sample frequency of 1000. Density curves and the ESS (Effective Sample Size) from the MrBayes output was analyzed using Tracer v. 1.7 (Rambaut et al., 2018). Resulting trees were visualized and edited in FigTree v.1.4.3 (Rambaut, 2016).

### 2.7 Analysis using species tree estimation

Species trees were inferred with two coalescence-based programs using the ML gene trees generated with RAxML. We used ASTRAL-II v. 5.5.7 (Mirarab and Warnow, 2015; Sayyari and Mirarab, 2016) and calculated support with local posterior probability (LPP). An additional analysis was performed with STAR (Liu et al., 2009) as implemented in the STRAW webserver (http://bioinformatics.publichealth.uga.edu/SpeciesTreeAnalysis/index.php). Individual gene trees were rooted online using the STRAW webserver. We selected species of *Gilbertiodendron* to root trees in all analyses.

### 2.8 Estimation of gene tree concordance

We assessed species-tree and gene-tree conflict and concordance with an emphasis on the *Anthonotha* clade. We used the concatenated-based species tree obtained with RAxML with 74 % matrix occupancy and 100% taxon completeness, and we also examined gene-tree conflict using the ASTRAL-II reference species tree. Concordance was quantified using the pipeline PhyParts (https://bitbucket.org/blackrim/phyparts) (Smith et al., 2015) and visualized with the ETE3 Python toolkit (Huerta-Cepas et al., 2016) as implemented in the script PhyPartsPieCharts (https://github.com/mossmatters/MJPythonNotebooks). Both analyses were performed with all branches regardless of their support, and also excluding branches with low support (-s 0.7 filter).

### 2.9 Mapping characters, ancestral state reconstructions and correlation of character states

We studied seven floral traits pertaining to floral symmetry, petals, stamens and staminodes (Table 1). Character states for all taxa were scored and obtained from published taxonomic and morphological studies of the Detarioideae (Bruneau et al., 2014; Tucker, 2002b, 2002a) and from more detailed studies in *Anthonotha* (Breteler, 2010, 2008), *Isomacrolobium* (Breteler, 2011), *Englerodendron* (Breteler, 2006; van der Burgt et al., 2007), *Pseudomacrolobium mengei* Hauman (De Wildeman, 1925; INEAC, 1951; IRCB, 1952; Wilczek et al., 1952), *Gilbertiodendron* (de la Estrella et al., 2014; de la Estrella and Devesa, 2014a, 2014b), *Didelotia* (Oldeman, 1964; van der Burgt, 2016), *Oddoniodendron* (Banak and Breteler, 2004), *Librevillea klainei* (Pierre ex Harms) Hoyle (Tucker, 2000), *Berlinia* (Mackinder, 2001; Mackinder and Harris, 2006; Mackinder and Pennington, 2011) and *Isoberlinia* (Tucker, 2002a). Additional information was obtained from available images and voucher specimens (Table S2). Our sampling encompassed all the morphological diversity for these floral traits observed in the extant species of the Anthonotha clade. For the remaining genera the scoring was done for the species representing each genus. Characters were scored as binary states (Table S3) and ancestral state reconstructions were performed on the tree obtained from the RAxML analyses using parsimony and likelihood methods as implemented in Mesquite ver. 2.75 (Maddison and Maddison, 2015). Maximum likelihood reconstructions were obtained with the Mk1 model of trait evolution.

**Table 1.** List of the four floral traits studied and the scoring for the genera sampled of the Berlinia clade.

| Floral trait |  |  |  |  |
| --- | --- | --- | --- | --- |
|  | State 0 | State 1 | State 2 | State 3 |
| 1. Flower symmetry | Zygomorphic | Actinomorphic |  |  |
| 2. Petal number | 6 petals | 5 petals | 1-4 petals | Absent |
| 3. No. of petal types | 1 type | 2 types | 3 types |  |
| 4. Petal merism | Variable | Invariable |  |  |
| 5. Fertile stamens | > 3 | 3 | < 3 |  |
| 6. No. of staminoids | > 6 | 6 | < 6 |  |
| 7. Stamen merism | Variable | Invariable |  |  |

Members of the Anthonotha clade vary in petal, stamen and staminode number and certain species also display reduction or increase in the number of organs within whorls. To determine whether changes towards an increase or decrease of organ numbers were correlated in their evolution, we tested the correlation between changes: 1) in petal and stamen numbers, 2) petal and staminode number, 3) flower symmetry and intraspecific variation in organ numbers, and 4) intraspecific variation in petal and stamen numbers. The characters were coded as binary factors (see Table S3 for scoring details) and pairs of characters were compared between ML independence and dependence models, and with Bayesian MCMC comparing the discrete dependent and independent models. After each comparison, likelihood ratios and Bayes factors were compiled to determine significance of correlation. These analyses were performed with BayesTraits v3 (Pagel and Meade, 2006).

### 2.10 Morphological characterization of petal types and their identity based on micromorphology

To determine whether differences in petal morphology along the dorsoventral axis within the flower were associated with different petal identities, we analyzed the morphology of each petal within each species using the same voucher specimens and source of information used for mapping characters noted above. Two parameters were used to establish the identity of each petal based on these arrangement types: the overall morphology of the petal (size, symmetry, and shape) and its location within the dorsoventral axis of the flower (adaxial, lateral or abaxial). This analysis focused on species of the *Anthonotha* clade where we were able to sample the entire extant diversity. Once the identity at the macro level was established for each species, we then examined whether all petals (adaxial, lateral and abaxial) have similar or different epidermal types, the latter suggesting differences in petal identity within the flower. We analyzed in total 21 species for the four genera *(Anthonotha, Isomacrolobium, Englerodendron* and *Pseudomacrolobium)* of our focal group (Table S4). Flowers from voucher specimens were first re-hydrated and later preserved in 70% ethanol. Petal micromorphology for each individual petal within each species was analysed on both sides with a light microscope (Nikon optiphot-2) and some specimens were further examined using a Zeiss scanning electron microscope (SEM) at high vacuum (EHT=5.00 kv). Classification of epidermal types and their level of differentiation were based on cell-shape traits (primary sculpture) and on the fine relief of the cell wall, or secondary sculpture (Barthlott, 1990). The terminology of these epidermal types follows (Ojeda et al., 2009).

## 3. Results

### 3.1 Bait capture and mapping success

We obtained an average of 919,856.5 reads per sample and recovered on average 84.89 % (± 3.94) of the baits in the set of samples included. Of these, an average of 35.77 % (± 13.15) of the reads mapped to the *A. fragans* reference contigs (Table S5). Overall, we obtained a coverage between 10-100X for at least 50% of the captured bait (Fig. S1). The final concatenated matrix consisted of 61 taxa, 922 exons (clusters) and 239,334 aligned bp with 74.06 % overall matrix occupancy (Table S6). The matrix contained 31,950 (13.35%) variable sites and 14,448 (6.04%) potentially parsimony informative sites. The final alignment is deposited on the Dryad Digital Repository: XXX.

### 3.2 Phylogenetic relationships within the Anthonotha clade

We obtained congruent topologies with ML (Fig. S2) and Bayesian (Fig. S3) analyses using the concatenated matrix, which resulted in resolved relationships and high support for all the lineages. The same lineages and topology were also recovered on the ML analysis using the individual genes (Fig. S4) and similar topologies were also obtained with the coalescent approaches using ASTRAL-II (Figs. S5) and STAR (Fig. S6). We did not obtain different topologies when only the 247 genes with the highest level of taxon completeness were included in the coalescent analyses of ASTRAL-II and STAR. We only noted a reduction in support of the relationship between *Anthonotha* ss and the cluster containing *Englerodendron, Isomacrolobium* and *Pseudomacrolobium,* and the remaining outgroup taxa (Fig. S5B). All these analyses identified the clade comprising *Englerodendron korupense, E. gabunense* and *Isomacrolobium graciflorum* with the lowest level of support relative to other lineages of our study (Figs. S2-S6). All the species included with multiple individuals and those with replicates formed sister clades, except *A. pynaerti, Englerodendron gabunense* and *E. korupense*. The analysis of gene congruence among the 247 genes with the highest level of taxon completeness revealed the presence of conflict of topologies with low frequency, and we did not observe discordant loci dominated by a single alternative topology. This discordance was concentrated in the Anthonotha clade, suggesting the presence of substantial incomplete lineage sorting in this group. Similar distribution of concordance and conflict was revealed using the ASTRAL-II species as a reference (Fig. S7) or the RAxML concatenated tree as a reference (Fig. S8). We did not find a difference in the level of conflict and concordance when only the highest supported branches were included (Fig. S7B and Fig. S8B).

In all our analyses, two main clades were recovered with high support within our focal group, one corresponding to all the *Anthonotha* ss species and the other containing the three remaining genera *Isomacrolobium, Englerodendron* and *Pseudomacrolobium* (Fig. 1). The latter monospecific genus has not been included before in phylogenetic analyses, and the grouping of these three genera was an unexpected result, given their diverse flower morphology (e.g. flower symmetry and size of petals).

### 3.3 Evolution of floral characters and ancestral state reconstructions

Our reconstructions suggest that the ancestral condition in the Anthonotha clade is a zygomorphic flower with five petals of two types, the largest on the adaxial side with a distinct morphology and the other four petals on the lateral and abaxial sides with a similar overall morphology. The ancestral reconstructions also suggest the presence of three large stamens on the adaxial side and six staminodes (Fig. 1H). This flower organization was observed in most species (12 out of 17) within *Anthonotha* ss. Our mapping and reconstruction analyses indicate that floral modifications consisting mainly in a reduction in number of petals, stamens and staminodes (Table 2 and Fig. S9), and more frequently in the *Isomacrolobium, Englerodendron* plus *Pseudomacrolobium* clade.

**Table 2.** Number of independent transitions inferred from the ancestral condition in the Anthonotha clade.

| Trait | Modification | Frequency |
| --- | --- | --- |
| Flower symmetry | Actinomorphy | 3 |
| Petal | Reduction of number | 6 |
|  | Increase of number | 1 |
|  | Reduction of petal types | 4 |
|  | Increase of petal types | 3 |
|  | Intraspecific petal number variation | 4 |
| Stamen | Increase of number | 1 |
|  | Reduction of number | 4 |
|  | Intraspecific stamen number variation | 3 |
| Staminoid | Reduction of number | 6 |

Our analyses suggest that the petal is the most labile organ among those we studied, involving modifications in petal types (morphology and location within the dorsoventral axis) (Fig. 2A) and number (organ merism) (Fig. 2B) with at least seven and eight independent transitions, respectively (Fig. S9). In addition to the seven modifications (increase and decrease) in petal types (Fig. 3A and C), we also identified three independent transitions to an alternative arrangement of two petal types. In these three independent transitions, the adaxial and the two lateral petals display the same morphology with the two abaxial petals having a distinctive morphology (Fig. 3B).

**Fig. 2.**
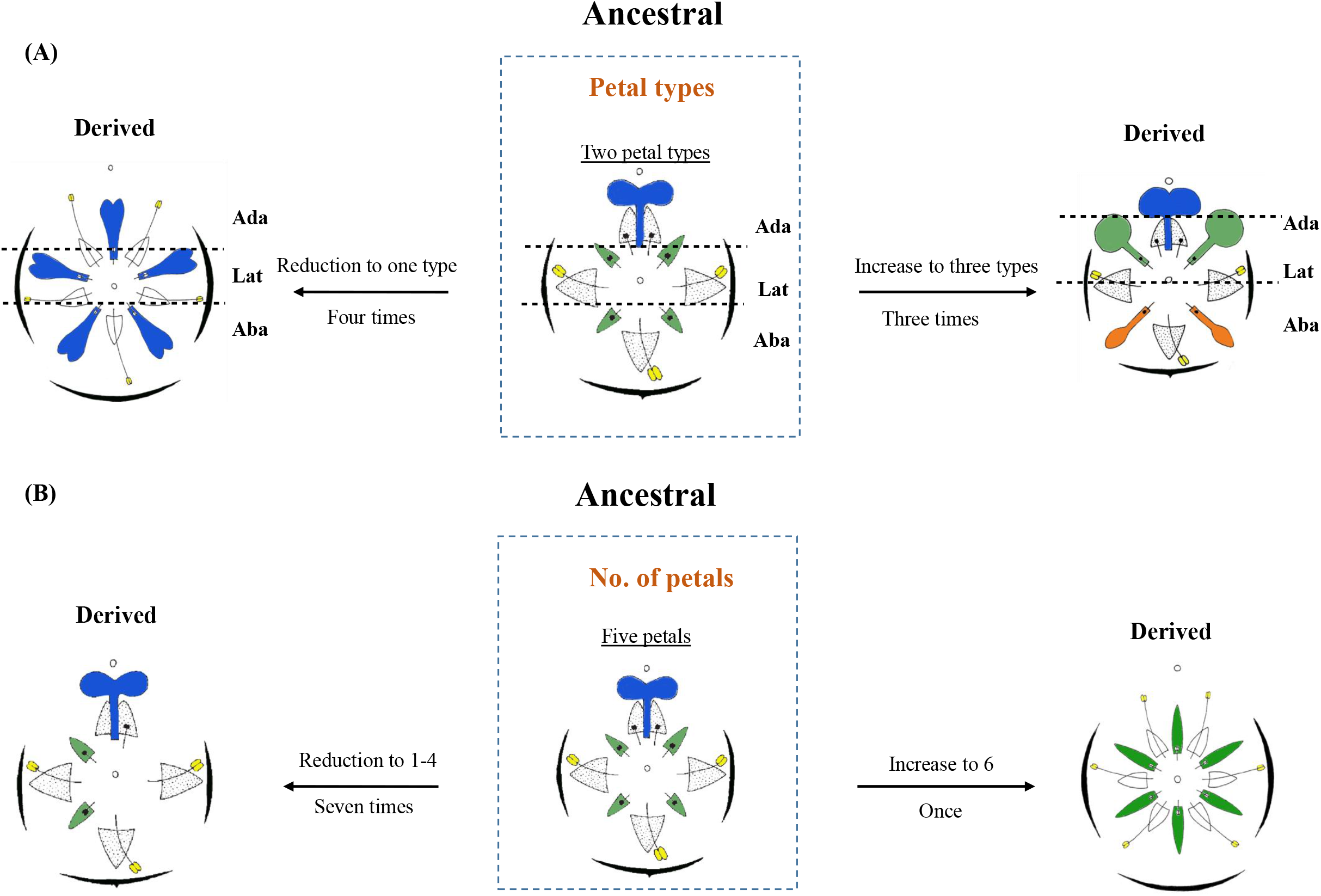
Schematic representation of the various modifications in petal types (A) and the number of petals (B) from the ancestral state reconstructed in the Anthonotha clade. Next to the arrows is indicated the type of transition and the number of independent transitions. Ada = adaxial petal, Lat = lateral petal, Aba = abaxial petal. Petal types (identity based on their position within the dorsoventral axis within the flower) are colored blue (adaxial), green (lateral) and orange (abaxial). Stamens are indicated in yellow and staminodes in black. Drawings modified fig. 3A2 in Breteler (2008: 141), fig. 15 in Breteler (2010: 84). Copyright Meise Botanic Garden and Royal Botanical Society of Belgium.

**Fig. 3.**
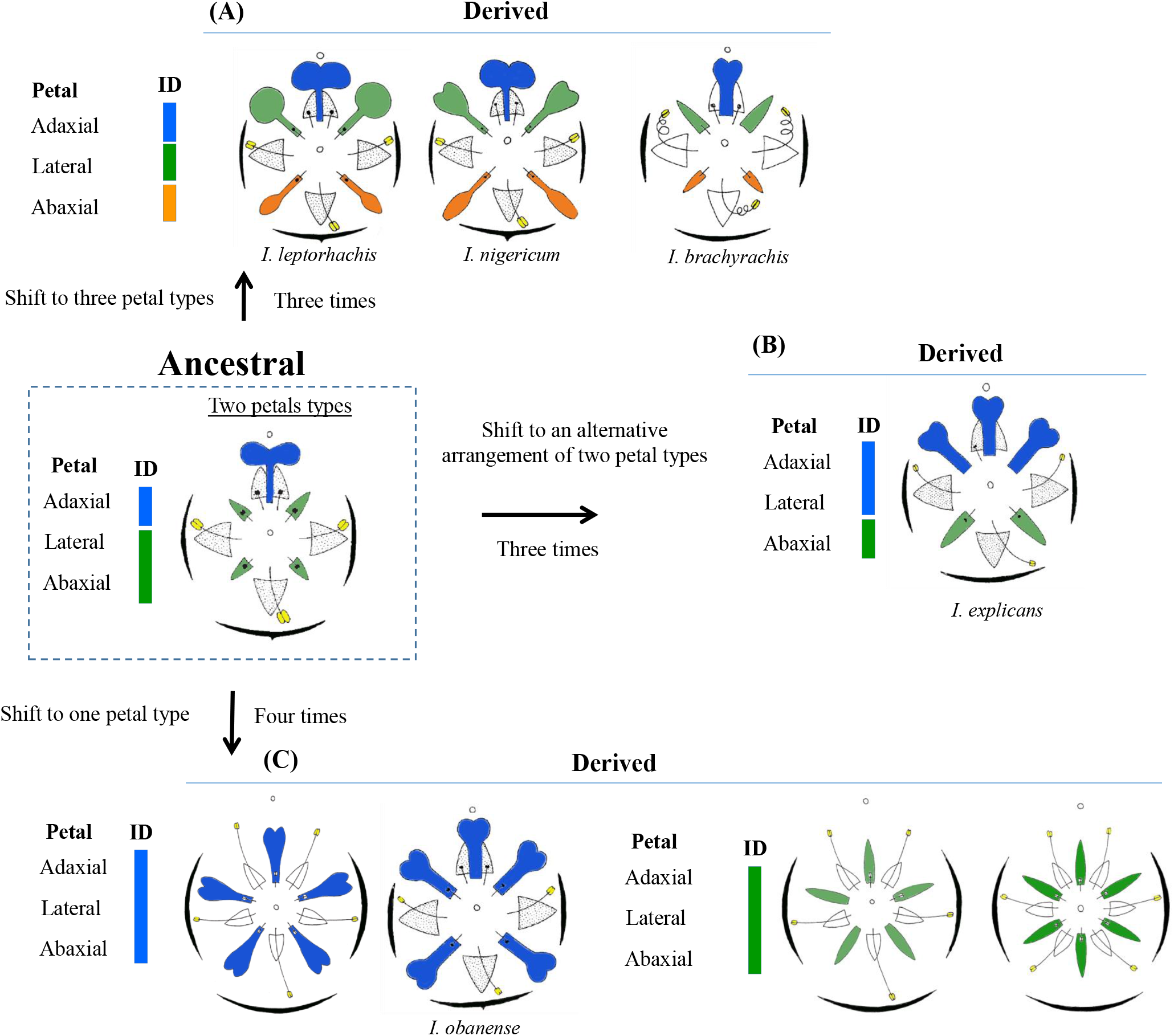
Schematic representation of the transitions in petal types from the ancestral state in the Anthonotha clade. Different colors represent petal identity (ID) based on their location along the dorsoventral axis within the flower and on their morphology on the corresponding diagrams: blue adaxial identity, green lateral identity and orange abaxial identity. Drawings modified from fig. 3A2 in Breteler (2008: 141), fig. 15 in Breteler (2010: 84), Fig. 4H, 17D, fig. 19B, fig. 22 in Breteler (2011:68, 75, 76, 78). Copyright Meise Botanic Garden and Royal Botanical Society of Belgium.

### 3.4 Correlation of floral character traits in the Anthonotha clade

Despite an overall trend towards a reduction in the number of petals, stamens and staminodes within this group, there is a lack of evidence that modifications in these three organs are correlated within a species. We also did not find evidence that flower symmetry is associated with the evolution of intraspecific variability in petal or stamen number. In addition, we did not find evidence that intraspecific variation within the petal whorl is correlated with intraspecific variation within the stamen whorl. Thus, our results suggest that within this group all these seven flower characters seem to have evolved independently.

### 3.5 Petal identity based on their location along the dorsoventral axis of symmetry and their corresponding micromorphology

Here we analyzed a total of 21 species within the Anthonotha clade. Of these, 11 species were similar to the ancestral condition inferred for the Anthonotha clade while seven species exhibited four independent transitions towards a reduction to one petal type. In addition, we also studied two species representing two of the three independent transitions to an alternative arrangement of two petal types and one species representing one (of two) independent shifts towards an increase in petal types (Table S4). We recorded two major epidermal types among all species that were examined (Fig. 4 and Table S4). All species have papillose cells, either conical cells with striations (PCS) or knobby cells with a rugose surface (PKR). PCS cells are characterized by a circular shape with the striations directed towards the highest point of the cell (Fig. 4C-E). In contrast, the cell shape of PKR is less rounded, the base of the cell is bigger (and with a square shape), and the striations of the rugose surface is more evenly distributed on the cell surface (Fig. 4H-I). Our analysis and comparison of epidermal types of each distinctive petal types (based on their morphology) and their location along the dorsoventral axis (adaxial, lateral and abaxial) within the flower, reveals a lack of epidermal micromorphology differentiation corresponding to differences in petal identity (Fig. 4). This suggests that, at the micromorphological level, all petal types within each species have a similar petal identity, regardless of position or morphology. However, we found a trend towards the presence of less differentiated cells on species with flowers where the petals are smaller or less exposed (Fig. S10).

**Fig. 4.**
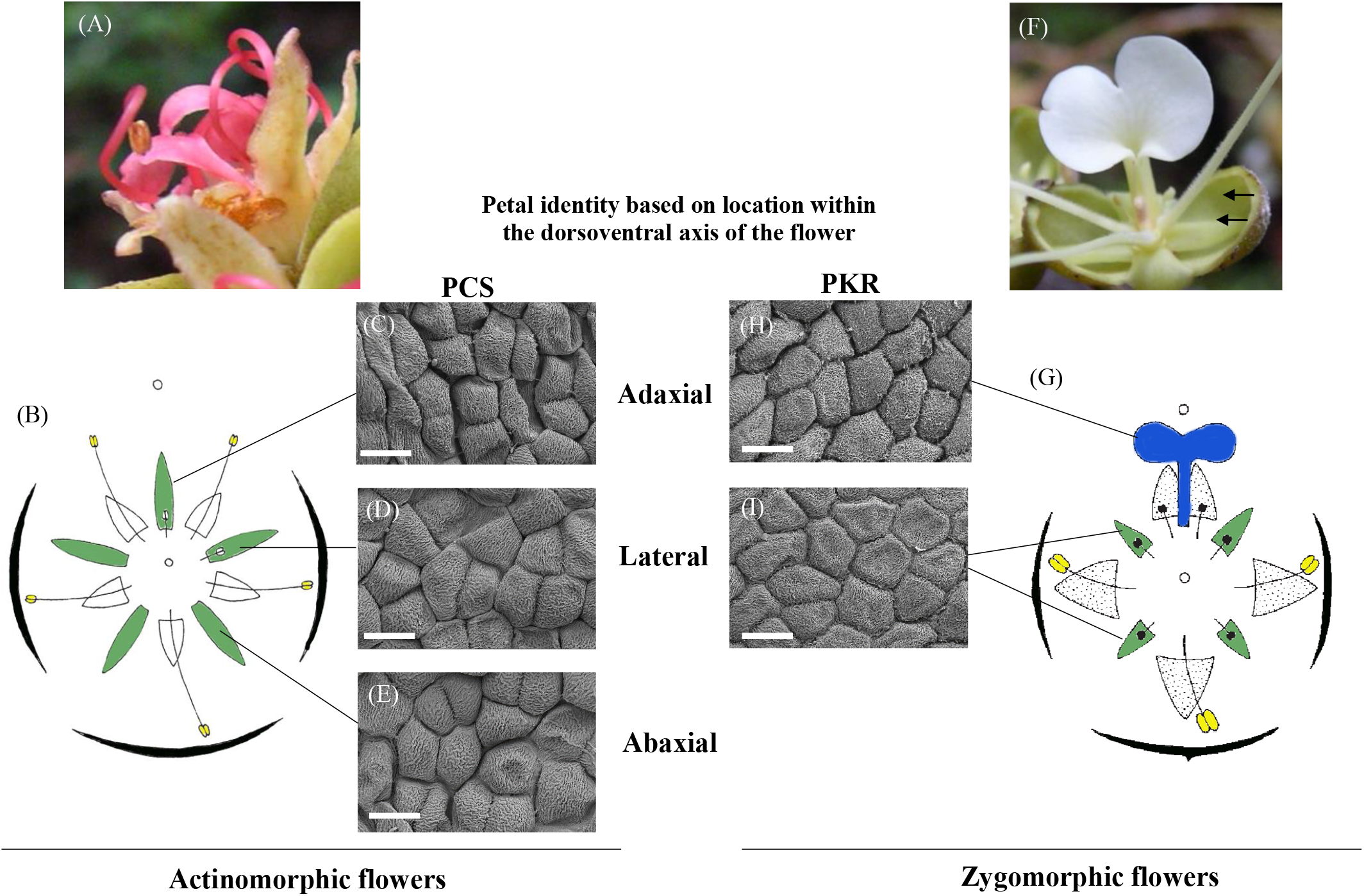
Distribution of epidermal types along the dorsoventral axis of symmetry within the flower. (A) Flower of *Englerodendron korupense* (XvdB_741_8) with all petals with similar petal macromorphology (one petal type), (B) schematic representation of *E. usambarense* indicating the location of adaxial, lateral and abaxial petals, (C-E) papillose conical cells (PCS) in a specimen of *E. conchylophlorum (Michelson 1060)* on three different petals selected based on their location in the flower. (F) Flower of *Anthonotha xanderi* (XvdB_279) with one large petal (adaxial) and lateral and abaxial petals of similar sizes (arrows), (G) schematic representation of *A. xanderi* with the identity of their petals based on their location. (H-I) papillose knobby rugose (PKR) cells observed in *Anthonotha noldeae (Carvahlo 6946)* on the two types of petals based on morphology. All scale bars 10 μm. Drawings modified from fig. 3A2 in Breteler (2008: 141), fig. 15 in Breteler (2010: 84). Copyright Meise Botanic Garden and Royal Botanical Society of Belgium. Photo credits: (A) Xander van der Burgt and (F) Ehoarn Bidault.

## 4. Discussion

### 4.1 The application of target enrichment as an integrative strategy to increase resolution at multiple taxonomic levels

Target enrichment (exome capture) is emerging as an efficient strategy for phylogenetic reconstruction across different taxonomic levels (Budenhagen et al., 2016; Mandel et al., 2014), allowing the combination of different studies using a common set of markers. Target enrichment is a cost-effective strategy to obtain markers for phylogenetic analysis at multiple taxonomic levels, including a set of markers for the entire flowering plants (angiosperms) (Budenhagen et al., 2016; Johnson et al., 2018). This approach has been successfully applied in several plant families (Carlsen et al., 2018; Comer et al., 2016; Herrando-Morairaa et al., 2018; Mandel et al., 2015, 2014; Moore et al., 2018) and it has proved useful to reconstruct relationships among closely related genera, within the same genus (Bogarin et al., 2018; Fragoso-Martíneza et al., 2017; Heyduk et al., 2016; Mitchell et al., 2017; Schmickl et al., 2016) and at species level (Nicholls et al., 2015; Villaverde et al., 2018). Within Leguminosae, this approach has been applied on lineages of the Papilionoideae and Caesalpinoideae (De Sousa et al., 2014; Nicholls et al., 2015; Ogutcen et al., 2018; Vatanparast et al., 2018) and our study is the first to use target enrichment outside these subfamilies using a complete sampling representing the entire extant diversity of the target group.

The recovery of the captured genes in target enrichment analyses have employed several strategies (or pipelines). PHYLUCE was developed for ultraconserved elements (UCEs) and uses a stringent filter to exclude paralogous sequences after the *de novo* assembly (Faircloth, 2015). HybPiper is also a common strategy used to recover captured regions. This strategy first maps the reads to a reference sequence and subsequently performs a *de novo* assembly, allowing the recovery of both exonic and intronic regions (Johnson et al., 2016). This pipeline identifies potential paralogous regions, but it lacks a specific guideline on how to handle paralogs after their identification. More recently, Fer and Schmickl (2018) developed HybPhyloMarker, which contains a set of scripts suitable from read quality to the reconstruction of phylogenetic trees; however, the identification and exclusion of paralogous is not specifically addressed. Unlike these bioinformatic pipelines, we employed an approach designed to recover orthologues from transcriptomes and low coverage genomes, which allows a more in depth assessment of paralogous sequences (Yang and Smith, 2014). This pipeline uses gene tree-based orthology approaches to identify paralogues regions (clusters) in the data set. This pipeline contains four strategies to identify and exclude these paralogous regions, and in this study we used the 1to1 strategy, which identifies homologues among the samples analyzed based on a strictly one-to-one assessment. This particular pipeline has been used both to select genes for the development of target enrichment (Vatanparast et al., 2018), as well as to recover captured regions for phylogenetic analyses (Nicholls et al., 2015).

Using this 1to1 approach, we were able to recover 67 *%* of the total bait size after quality filtering and this data set allowed us to increase the level of resolution among closely related genera, providing strong phylogenetic support when applied to all extant species within *Anthonotha* ss (Fig. 1). This also allowed us to infer relationships within this genus and the other three closely related genera, despite of the potential for incomplete lineage sorting (ILS) observed in our data sets (Figs. S7-S8). Our preliminary analyses at higher taxonomic levels (including nearly all described genera within Detarioideae) suggested the Detarioideae bait set and the strategy used to recover orthologues is suitable for reconstructing highly supported relationships at the subfamily level (M. de la Estrella *et al.,* in prep.). This hybrid capture approach will increase the level of resolution compared to previous analyses (Bruneau et al., 2008; de la Estrella et al., 2017; LPWG, 2017) and allow the possibility to build a more comprehensive phylogeny of the subfamily at multiple taxonomic levels. This strategy is also suitable for analyses at the intraspecific level, as the application of this bait set has recovered enough signal to reconstruct the population structure and demography of *Anthonotha macrophylla,* one of the most widely distributed species in the Guineo-Congolian region (S. Cervantes *et al.,* in prep.).

### 4.2 The Anthonotha clade exhibits remarkable floral diversity at multiple levels with overall tendency to reduction in organ number

Detarioideae and Dialioideae are renowned for their high floral diversity, including floral symmetry, and shifts in number and fusion of organs. Although this diversity and level of variation have been documented in major clades (Bruneau et al., 2014; Zimmerman et al., 2017), it has rarely been studied in detail among closely related genera (or within a genus) using a complete taxon sampling and a well resolved phylogenetic framework. One of our major findings is the extensive diversity in the number of transitions within the Anthonotha clade, with at least 35 instances of modifications in the seven floral traits we analyzed, suggesting that at least 30% of the species have modified at least one of these floral traits (Table 2). Our analyses also suggests that this group has an overall tendency towards a reduction of floral organs, and in particular a reduction in the number of petals (Table 2 and Fig. 2). Petal number reduction, however, occurs only with the lateral and abaxial petals, while the adaxial petal is always retained, except in the most extreme condition of complete lack of petals. Apetalous flowers are commonly observed in other Detarioideae genera and are frequent in the resin-producing clade (Fougère-Danezan et al., 2010), where apetaly is constant within a genus. In Detarioideae, 20% (17 of 81) of genera are considered apetalous and this lack of petals is observed in almost all species within these genera (Bruneau et al., 2014; de la Estrella et al., 2018; Tucker, 2000). This is the case, for example in *Didelotia* (Fig. 1B), a genus of 11 species where all species are apetalous. In our focal group, only one species, *Isomacrolobium vignei,* has been reported with a complete absence of petals, and this is observed in only some of the specimens analyzed (Breteler, 2011), thus highlighting the remarkable lability of this trait at multiple levels within the Anthonotha clade.

In this study we found that the petal is the most labile of the floral traits that we studied. Flowers of the four genera in the Anthonotha clade display high diversity of petal sizes, number, shape and arrangement in the flower (Figs. 2 and 3). Some species, such as *Isomacrolobium leptorhachis, I. nigericum* and *I. brachyrachis,* display different petal shapes and sizes along the dorsoventral axis of the flower, similar to some extent to the level of morphological differentiation of the papilionoid flower, with one adaxial (dorsal), two lateral, and two abaxial (ventral) petals. A different arrangement is observed in the four *Englerodendron* species with actinomorphic flowers (Fig. 3C), where all petals display the same size and morphology within the same flower, but considerable diversity in shape and size is observed among the four species. One species (E. *conchyliophorum)* has all petals resembling the large adaxial (dorsal petal) typical of the zygomorphic flowers of most of the genera in the Berlinia clade (Bruneau et al., 2014), while in the remaining three species all petals resemble those from lateral petals (Breteler, 2006; van der Burgt et al., 2007).

The Anthonotha clade also displays high levels of intraspecific variation in petal and stamen number with four and three independent transitions, respectively (Table S2). A high level of intraspecific variation has been observed in early diverging angiosperms, including lineages of eudicots sister to core asterid and rosid clades (Ronse De Craene, 2015). However, intraspecific variation is not a common feature of the more derived core eudicot clades. In our focal group, the most extreme level of variation has been reported in *Isomacrolobium vignei,* with intra-individual variation in the number of petals ranging from 0 to 4 (Breteler, 2011). Such levels of variation have been reported in a few instances in core eudicots (Kitazawa and Fujimoto, 2014); for instance in *Cardamine hirsuta* (Brassicaceae) where the intra-individual variation in petal number is due to loss of developmental robustness and the evolution of a selfing syndrome (Monniaux et al., 2016). However, the mechanisms behind this variability is not clear in the case of *I* *vignei* and further work is required on this species.

### 4.3 A different molecular mechanism behind petal identity in the Anthonotha clade and Detarioideae?

One of the main questions arising from our analyses is what is maintaining this diversity of floral traits among closely related taxa. Tucker, (2002a, 2002b) has characterized the developmental changes that occur in taxa that display modifications in petal number in a number of Detarioideae. However, despite recent advances in the understanding of the molecular basis of petal identity and petal symmetry in some model Papilionoideae species *(Lotus* and *Pisum*), it is not clear whether the same transcription factors *(CYCLOIDEA, WOX1, MIXTA-like* and *MADS-box)* (Feng et al., 2006; Wang et al., 2010, 2008; Weng et al., 2011; Xu et al., 2013; Zhuang et al., 2012) are also implicated in the evolution of the diverse morphology observed in Detarioideae. Analyses of gene expression in several genera of the Papilionoideae have further demonstrated the role some of these transcription factors play during the evolution of petal morphology. For instance, the dorsalization (the acquisition of the morphology of the dorsal petal) of lateral and ventral petals in *Cadia purpurea* (G. Piccioli) Aiton is the result of a homeotic transformation due to an expansion of the expression domain to the lateral and ventral domains of the *CYCLOIDEA* gene conferring dorsal identity (Pennington et al., 2006). Similarly, the lateralization of the dorsal and ventral petals among *Lotus* species of the Macaronesia region have been associated with a shift in the timing of expression of the *CYCLOIDEA* gene responsible for lateral identity (Ojeda et al., 2017). Furthermore, specimens of *Lathyrus odoratus* L. with modified dorsal petals (hooded mutant, hhd) have been explained by alterations of gene expression and sequence truncation of the *CYCLOIDEA* dorsal identity gene, which resulted in a homeotic alteration resulting in the lateralization of the dorsal petal in this loss of function mutant (Woollacott and Cronk, 2018). In all these species, modifications in petal identity are also accompanied by modifications in the petal micromorphology associated with each identity. As such, in the papilionoid flower the typical petal micromorphology characteristic of each petal along the dorsoventral axis has been altered during changes in petal morphology (Ojeda et al., 2009).

In the Anthonotha clade, although we observed epidermal types similar to those previously reported in other legumes (Ojeda et al., 2009), their distribution on the petal types along the dorsoventral axis does not suggest that each petal has a different identity. Even on the species with three distinct petal types *(Isomacrolobium leptorhachis, I. nigericum,* and *I. brachyrachis)* with one adaxial (dorsal), two lateral and two abaxial (ventral) petals (Fig. 3 A), all five petals have the same petal micromorphology. This is congruent with previous analyses of more limited sampling outside the Papilionoideae, which suggested that the micromorphological differentiation along the dorsoventral axis of the flower is unique to this subfamily (Ojeda et al., 2009). We also did not find an association of a particular epidermal type with a specific petal type in species with zygomorphic flowers, but there is a tendency for PKR cells to occur more often (twice) than PCS cell types and the opposite trend is observed in the four species with actinomorphic flowers, with PCS recorded more often (three times) than PKR cell types (Table S4).

Among the four genera we studied in detail, *Isomacrolobium* has the highest level of diversity of petal types. In this genus only one species displays the ancestral state inferred for the Anthonotha clade and we observed three different modifications to the ancestral type arrangement. But even in this group there is no association with a specific epidermal type. For example, in *I. explicans*, where the lateral petals resemble the adaxial (dorsal) petal, suggesting the possibility of dorsalization of the ventral petals, there are no differences in epidermal morphology (Fig. 3B). We found the same for the only species of *Isomacrolobium* with zygomorphic flowers, *I. obanense,* where all petals resemble the adaxial (dorsal) petal (Fig. 3C).

Overall, our results provide a solid phylogenetic framework to further explore detailed comparisons among species with contrasting morphologies and to analyze in more detail the molecular basis underlying this diversity on petal morphology.

## Acknowledgements

We thank the staff for the Meise and Kew herbaria for their support during the visits and collection of material. M.E. was funded by the European Union’s Horizon 2020 research and innovation programme under the Marie Sklodowska-Curie grant agreement No 659152 (GLDAFRICA). This work was supported by the Fonds de la Recherche Scientifique-FNRS (F.R.S.-FNRS) under Grants n° T.0163.13 and J.0292.17F, and by the Belgian Federal Science Policy Office (BELSPO) through project AFRIFORD from the BRAIN program. Permission to reproduce photographs was generously given by C. Jongkind/Fauna & Flora International.

## Author Contribution

DIO and OH designed the study; DIO, EK, ME, SC, EB, JM, BD participated in the bait design and performed the laboratory analyses. DIO and EB conducted the analyses. DIO wrote the manuscript. SJ and BD contributed to collection of plant material. All authors read the first draft and provided comments.

## Supporting Information

### Figures

**Fig. S1.** Capture success of the baits on the 61 samples after mapping with the 1021 exons reference.

**Fig. S2.** Best tree recovered with RAxML on the concatenated matrix using 200 bootstrap replicates.

**Fig. S3.** Best tree recovered with MrBayes on the individual clusters with the Bayesian support.

**Fig. S4.** Best tree recovered with RAxML on individual clusters (orthologues) using 200 bootstrap replicates.

**Fig. S5.** ASTRAL species tree inference based on the ML inferred individual genes (clusters) obtained with RAxML and fast 200 boostrap support. (A) Using the longest 247 rooted trees (B) and using all 661 rooted trees. Branch support is indicated above each branch. Red represent the *Anthonotha* ss clad and blue *Englerodendron, Isomacrolobium* and *Pseudomacrolobium.*

**Fig. S6.** STAR species tree inference using the ML inferred individual genes (clusters) obtained with RAxML and 200 fast boostrap support. (A) Using the 247 more complete rooted trees and with (B) including all 661 rooted trees. Red represent the *Anthonotha* ss clad and blue *Englerodendron, Isomacrolobium* and *Pseudomacrolobium.*

**Fig. S7.** ASTRAL-II tree of the Anthonotha clade with summary of conflict and concordant gene trees with (A) including all branches in the analysis and (B) including only branches with high support (70%). Pie chart color coding: blue: fraction of gene trees supporting the shown split; green: fraction of gene trees supporting the second most common split; red: fraction of gene trees supporting all other alternative partitions; gray: fraction of gene trees with no information (missing or unresolved).

**Fig. S8.** RAxML concatenated tree of the Anthonotha clade with summary of conflict and concordant gene trees with (A) including all branches in the analysis and (B) including only branches with high support (70%). Pie chart color coding: blue: fraction of gene trees supporting the shown split; green: fraction of gene trees supporting the second most common split; red: fraction of gene trees supporting all other alternative partitions; gray: fraction of gene trees with no information (missing or unresolved).

**Fig. S9.** Ancestral state reconstruction of flower symmetry, petal numbers, intraspecific stamen variation, no. of stamens, no. of staminodes, petal types and no. of petals with maximum likelihood.

**Fig. S10.** Level of differentiation on papillose conical cells (PCS) on species with actinomorphic flowers in *Englerodendron.* (A) *E. korupense (van der Burgt 741)* with smaller petals and less differentiated PCS cells (B-E). (F) *E. conchylophlorum (Michelson 1060)* with larger petals (and more area exposed) with PCS showing a higher level of differentiation (G-J). All scale bars 10 μm. Drawings modified from fig. 3A2 in Breteler (2008: 141), fig. 15 in Breteler (2010: 84). Copyright Meise Botanic Garden and Royal Botanical Society of Belgium.

### Tables

**Table S1.** Assembly metrics from the four transcriptomes used to generate the Detarioideae bait.

**Table S2.** Voucher specimens and location of the species included in the analyses.

**Table S3.** Matrix of the character states used in the ancestral reconstruction in the Berlinia clade.

**Table S4.** Voucher specimens of species used in the analysis of petal micromorphology. All specimens collected from Meise herbarium. Epidermal types identified in the species analyzed. PCS = papillose conical cells with striations, PKR = papillose knobby rugose cells,^1^ = presence of trichomes.

**Table S5.** Statistics of the capture for the samples included in this study.

**Table S6.** Supermatrix dimension, statistics of the number of orthologues and percentage of success for the species included in the study.

