## Supplementary figures and images for "Phylogenomics within the Anthonotha clade (Detarioideae, Leguminosae) reveals a high diversity in floral trait shifts and a general trend towards organ number reduction"

### Fig S1

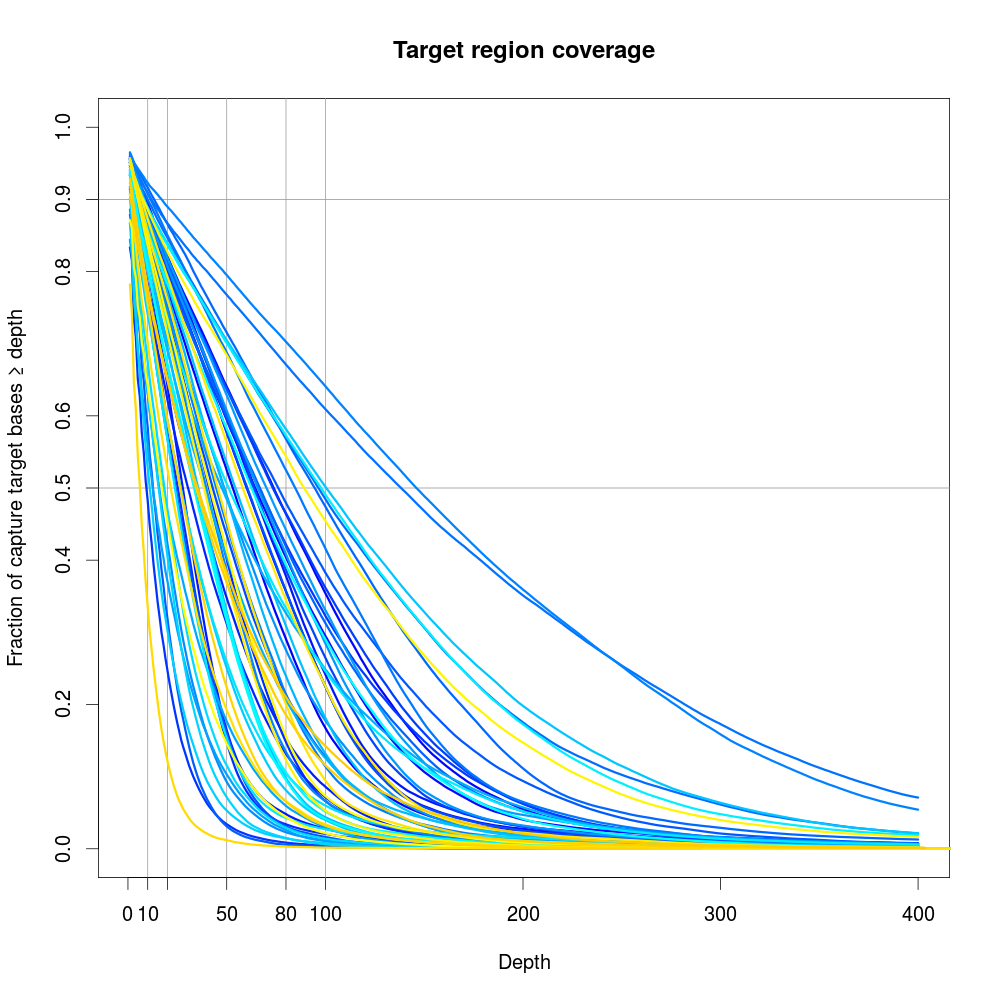

### Fig S2

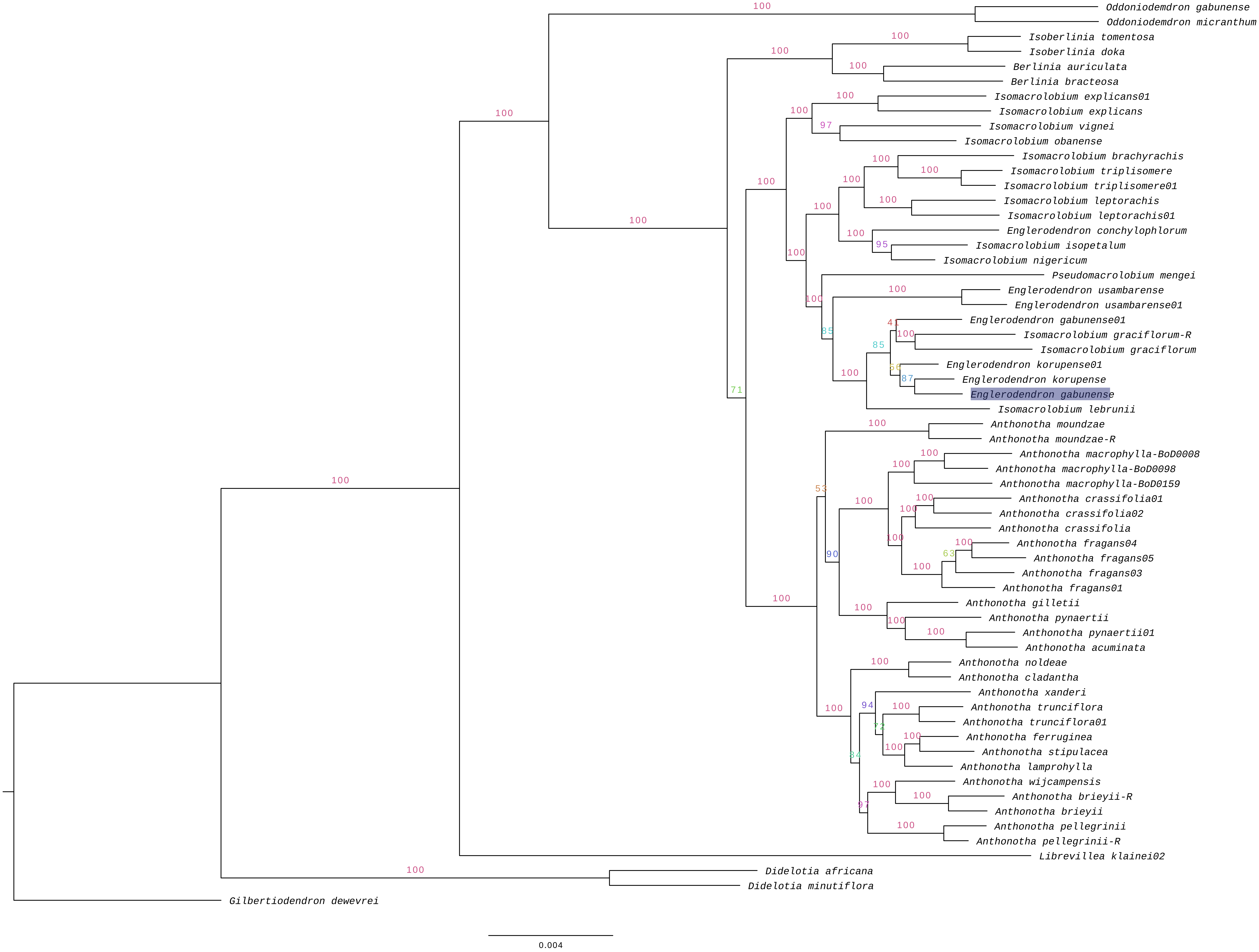

### Fig S3

#### Slide 1
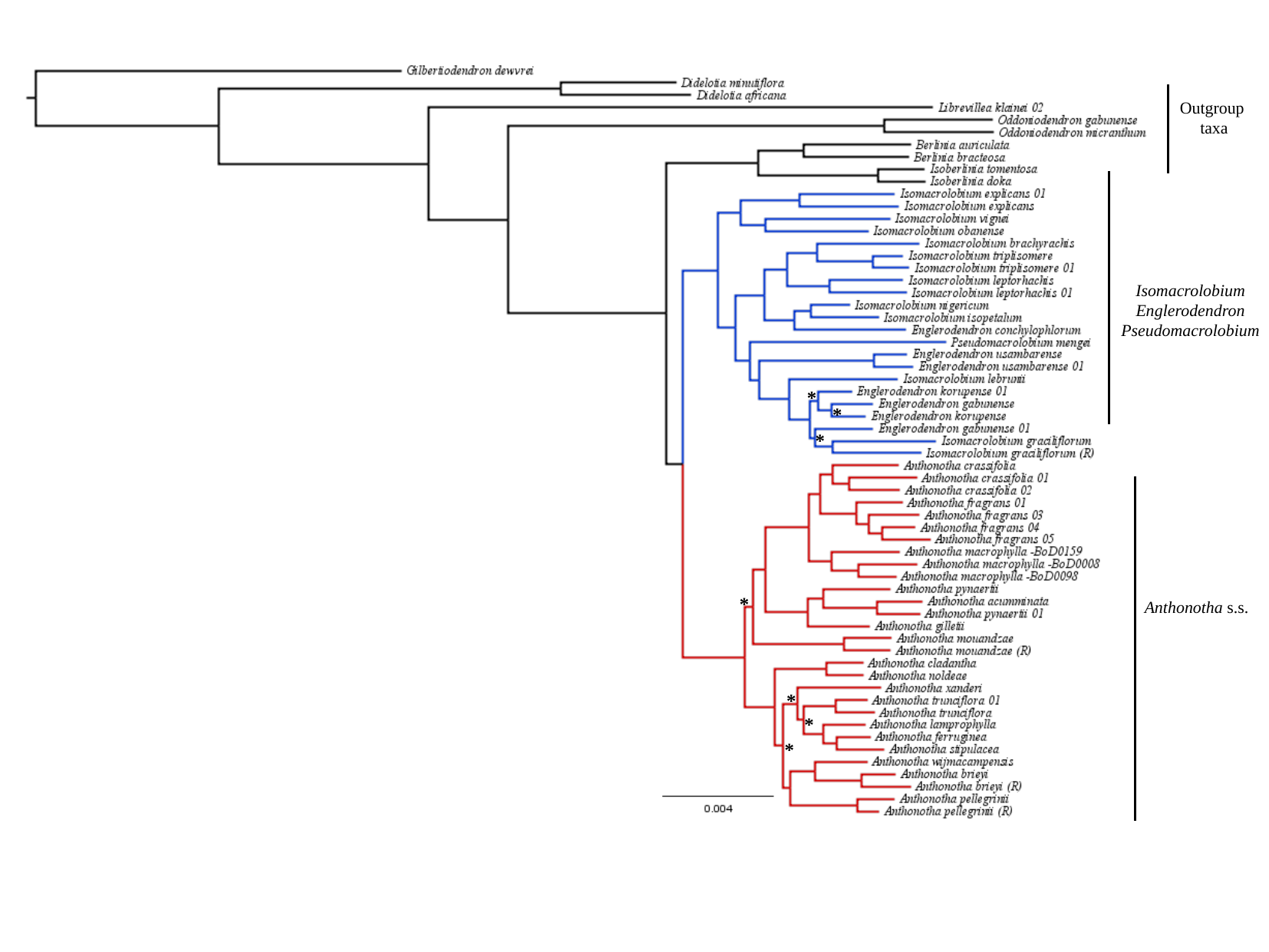

Outgroup
taxa
Isomacrolobium
Englerodendron
Pseudomacrolobium
*
*
*
*
Anthonotha s.s.
*
*
*

### Fig S4

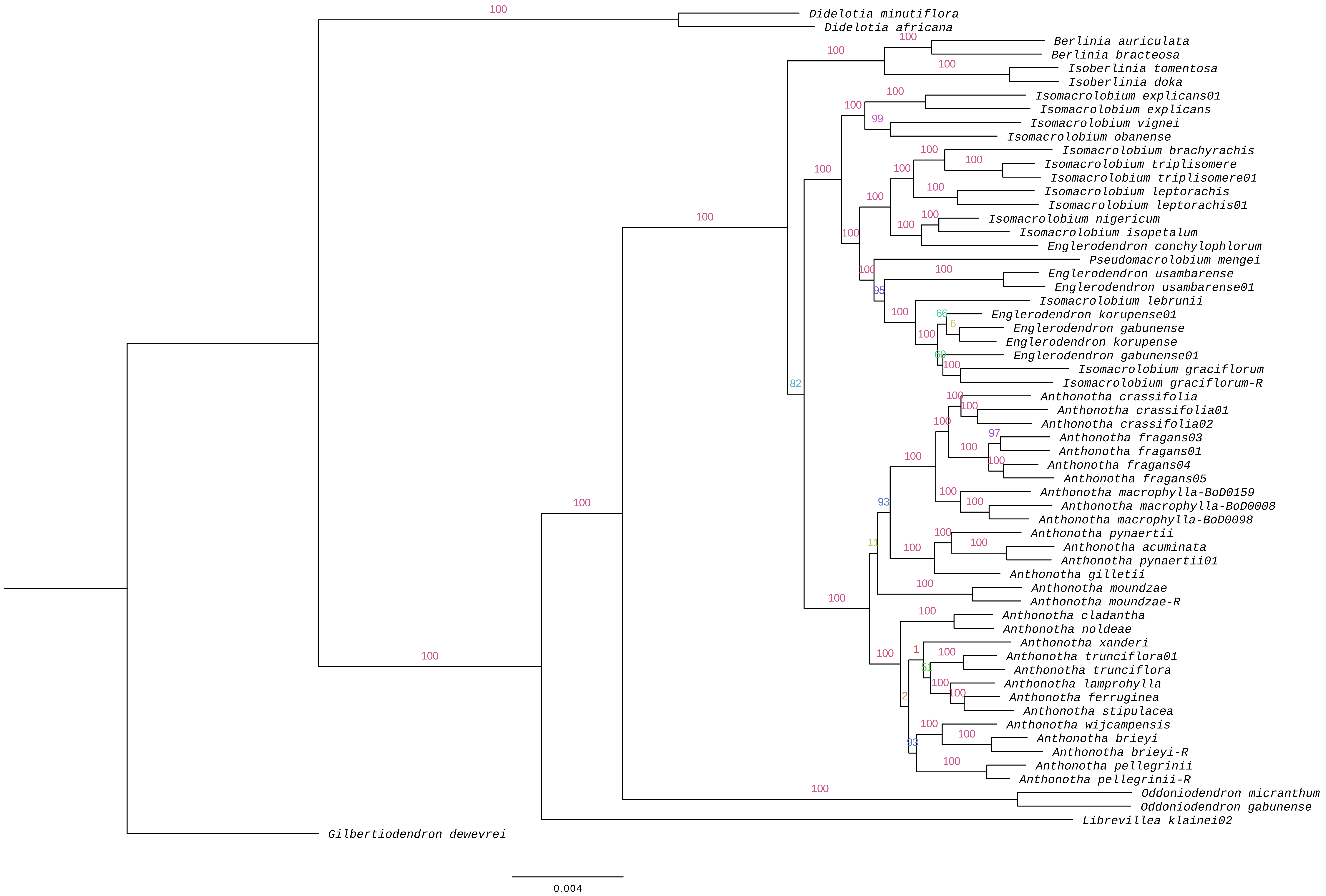

### Fig S5

#### Slide 1
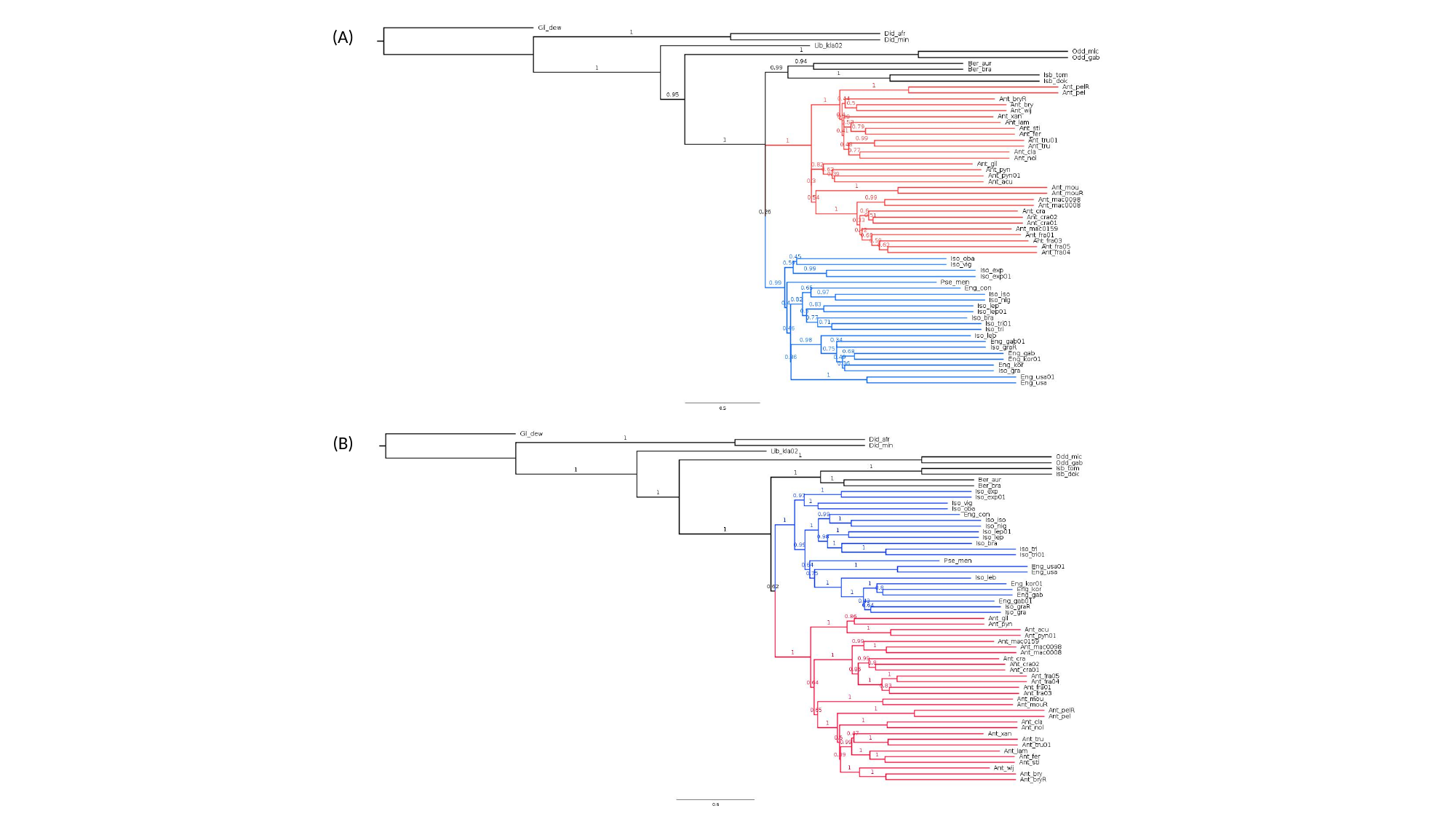

(A)
(B)

### Fig S6

#### Slide 1
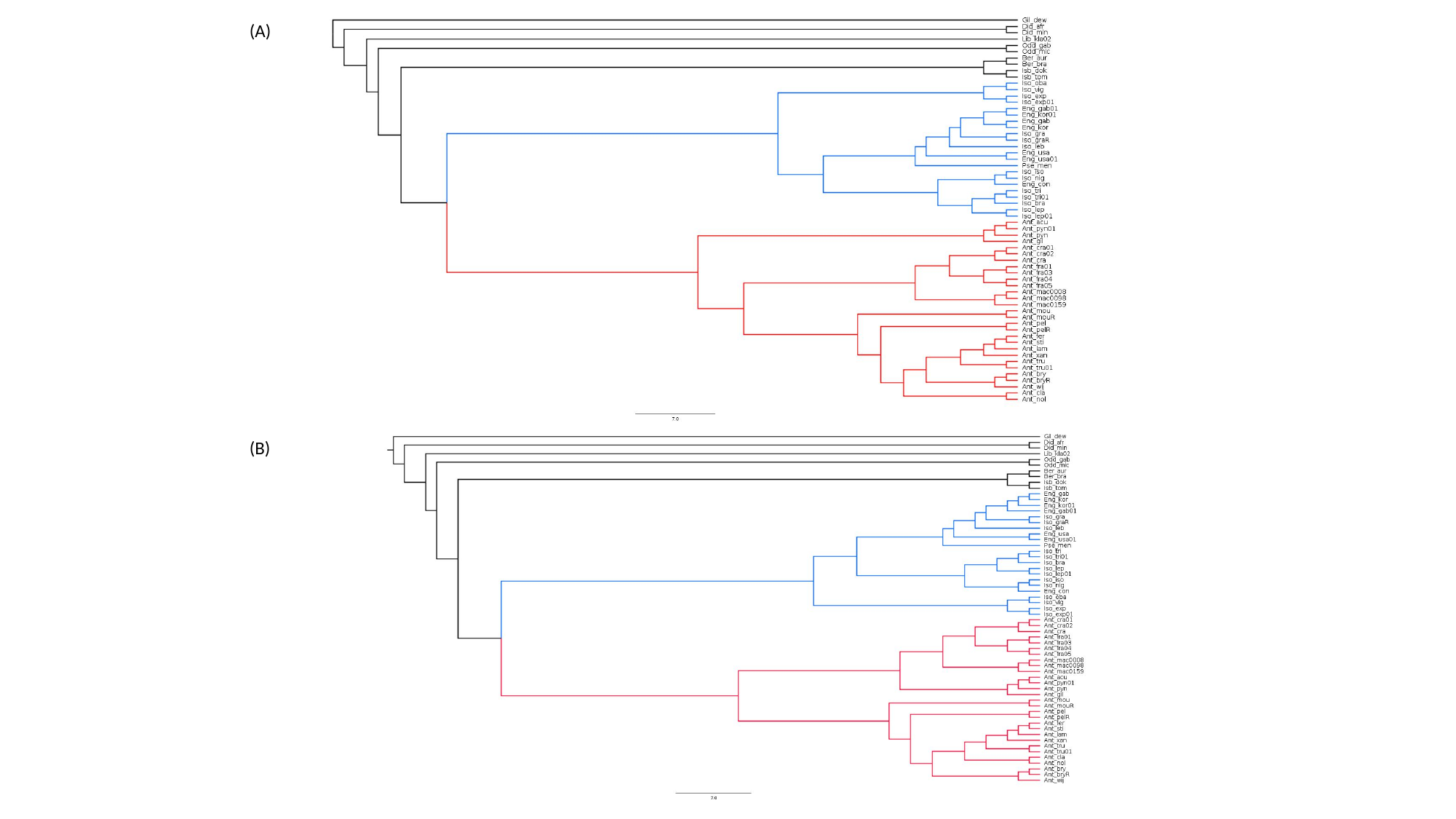

(A)
(B)

### Fig. S9

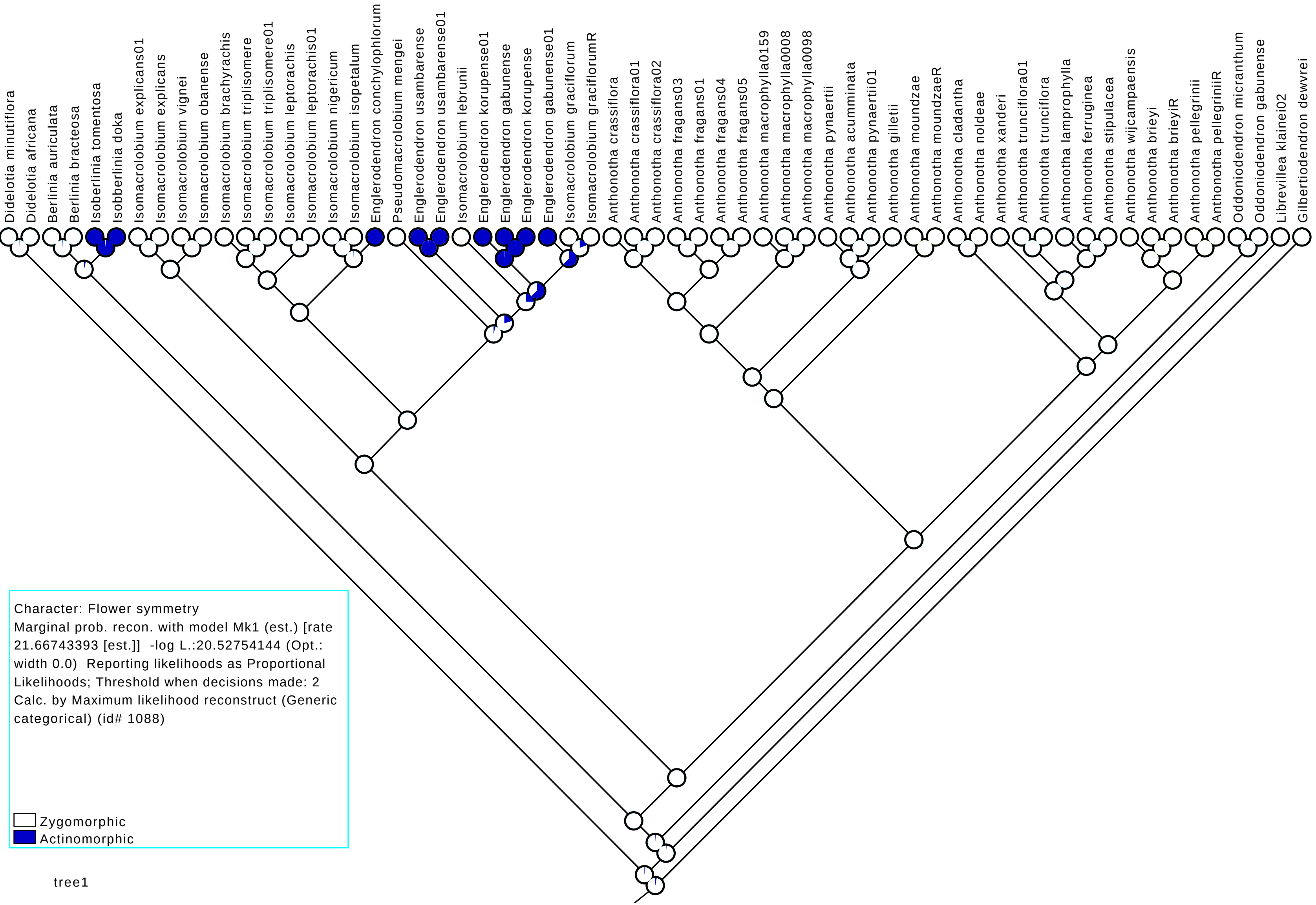

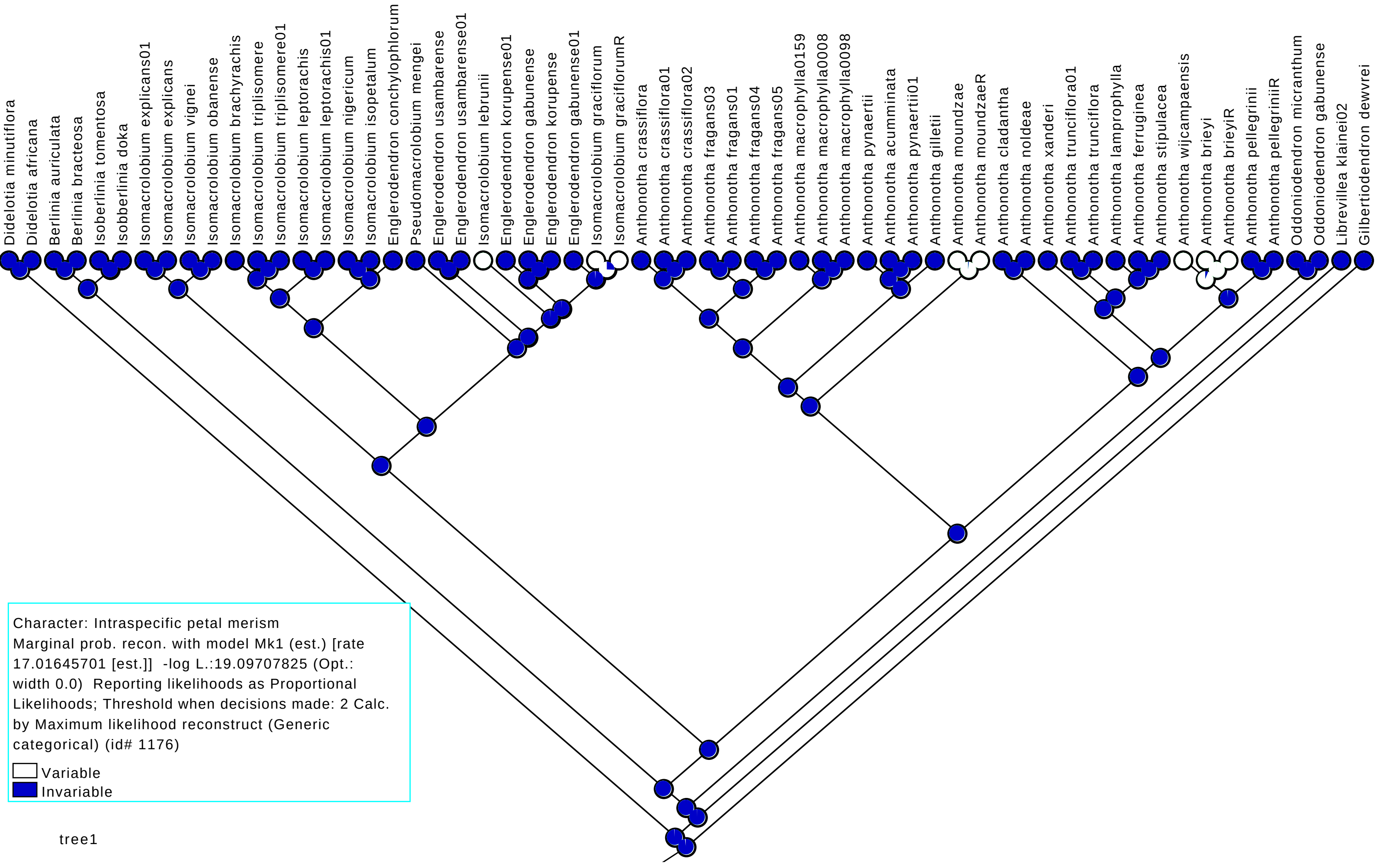

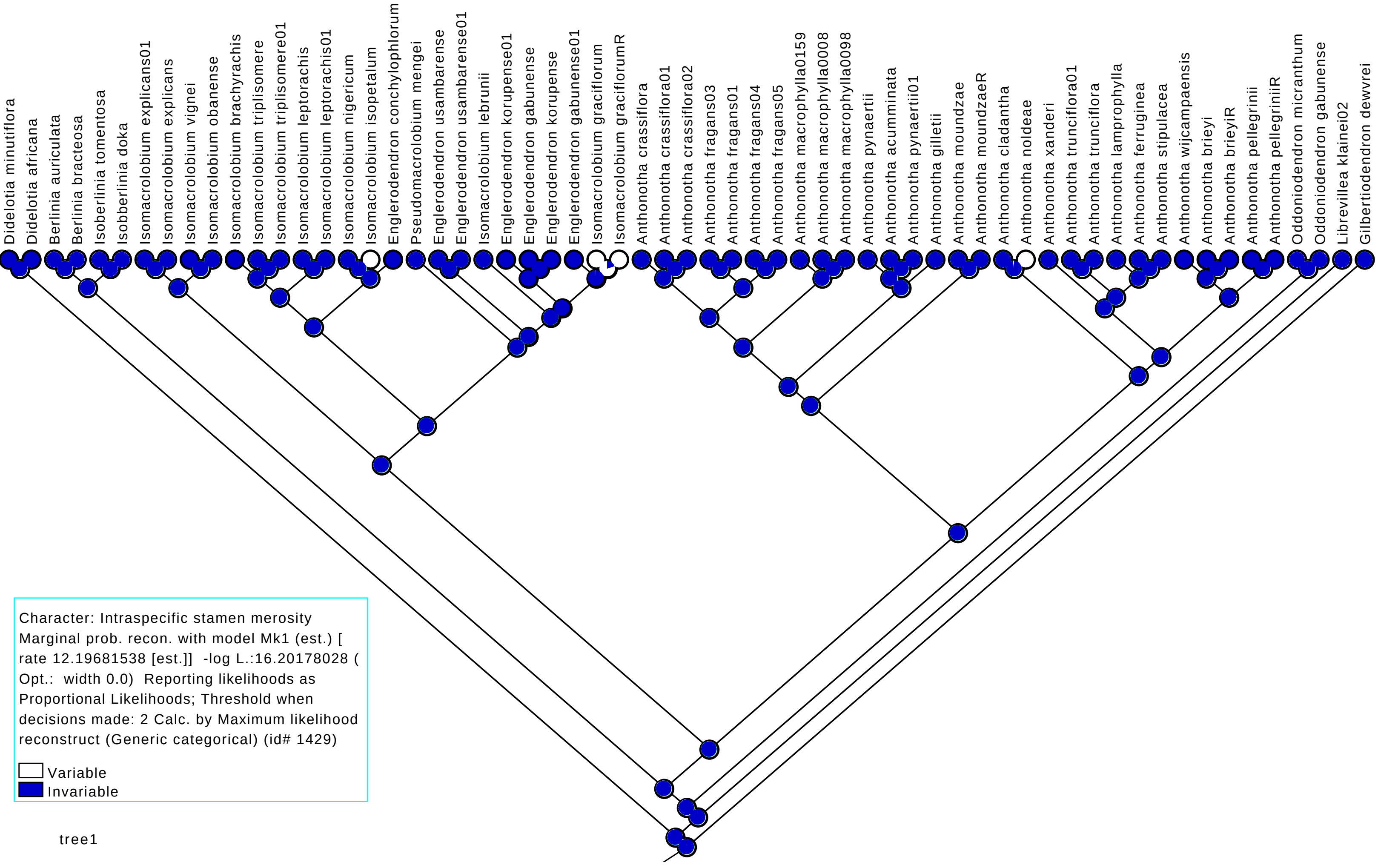

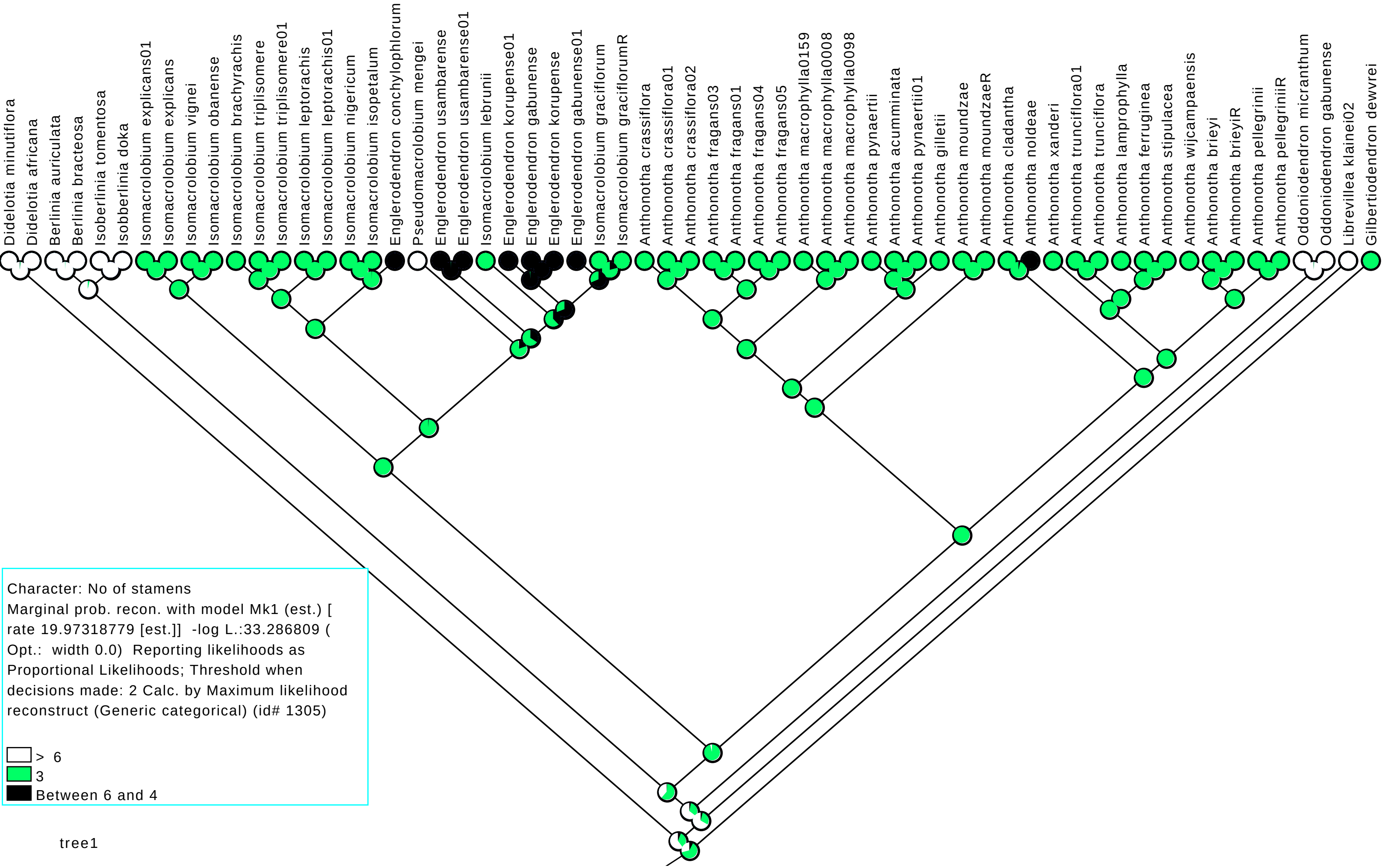

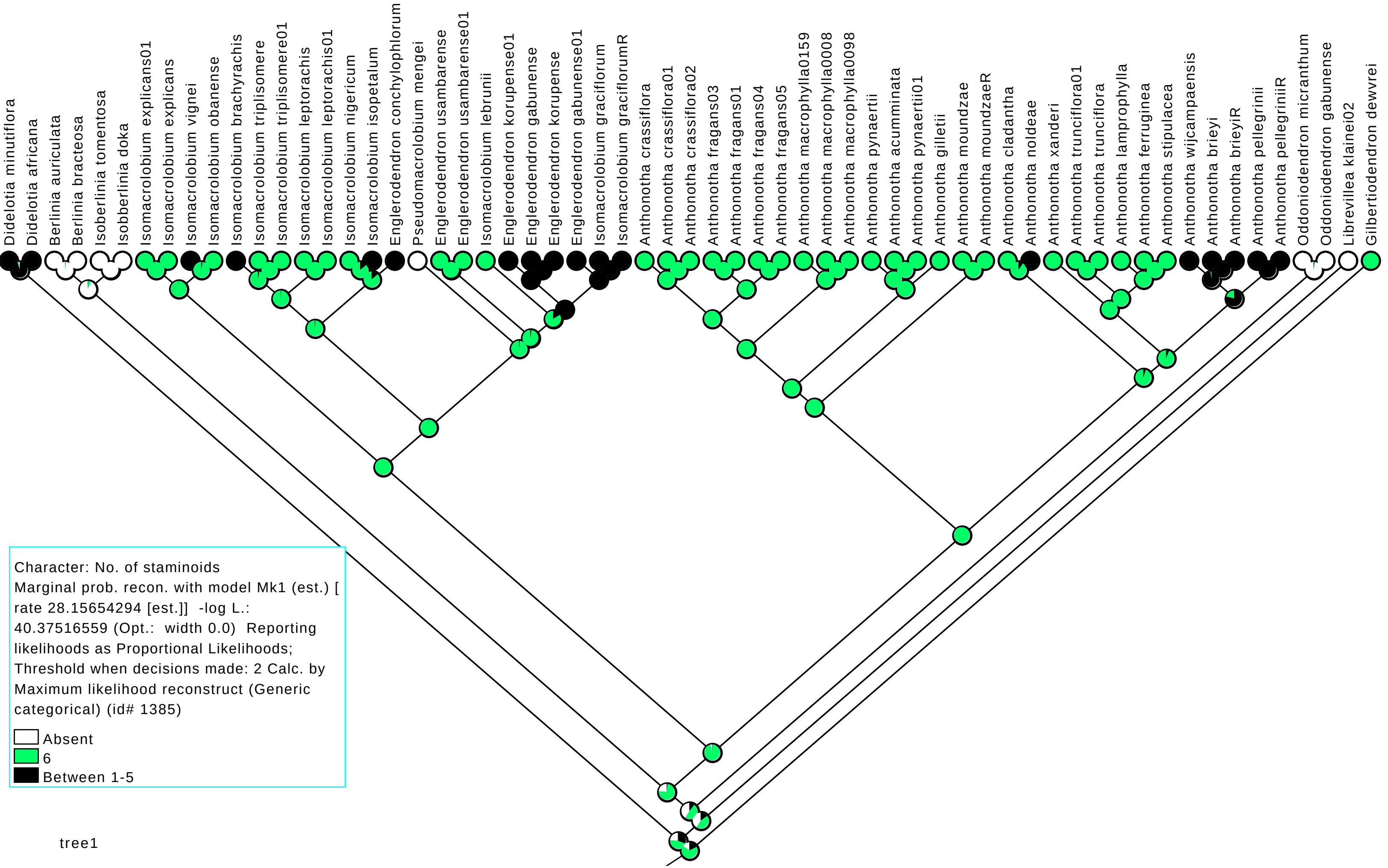

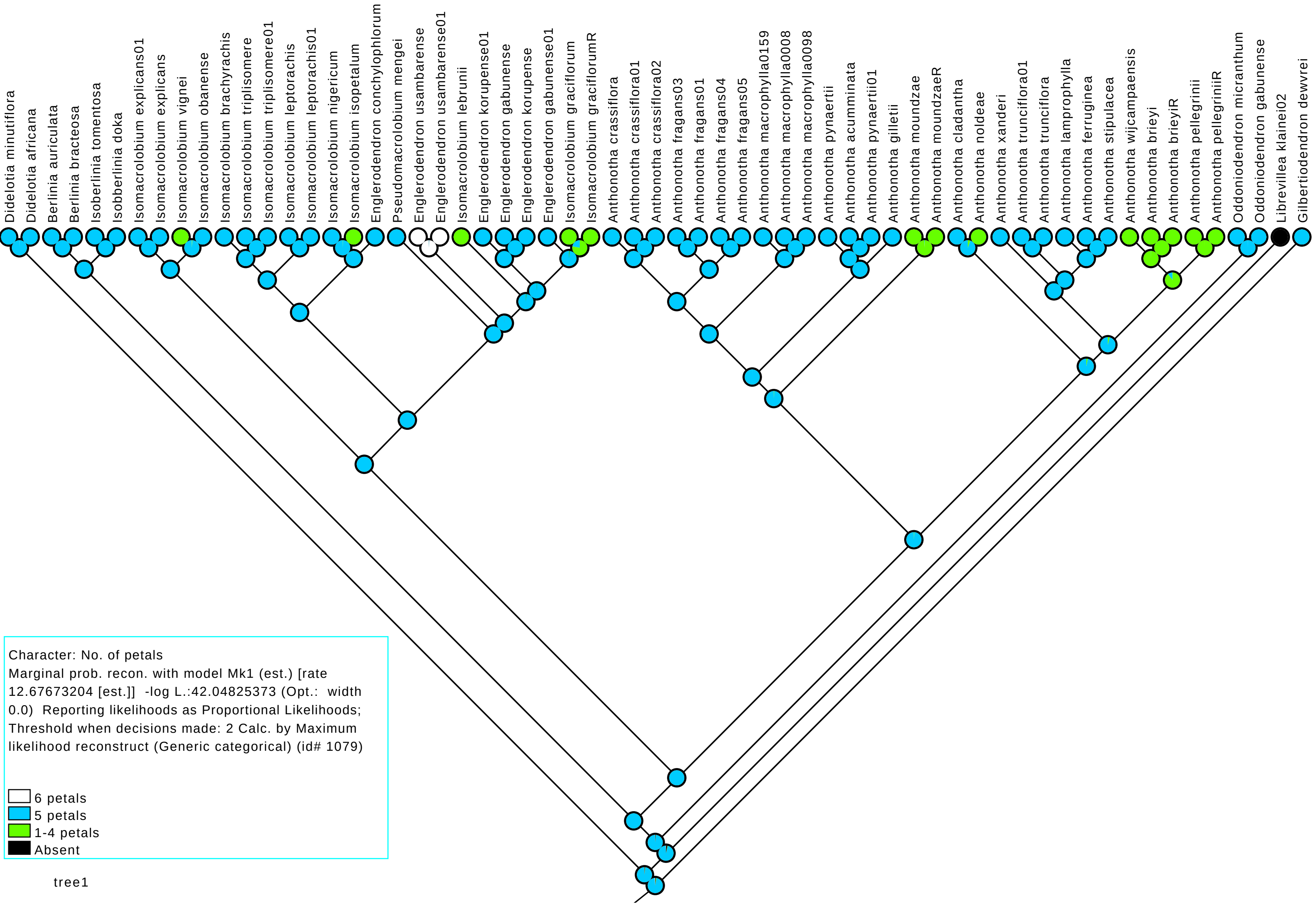

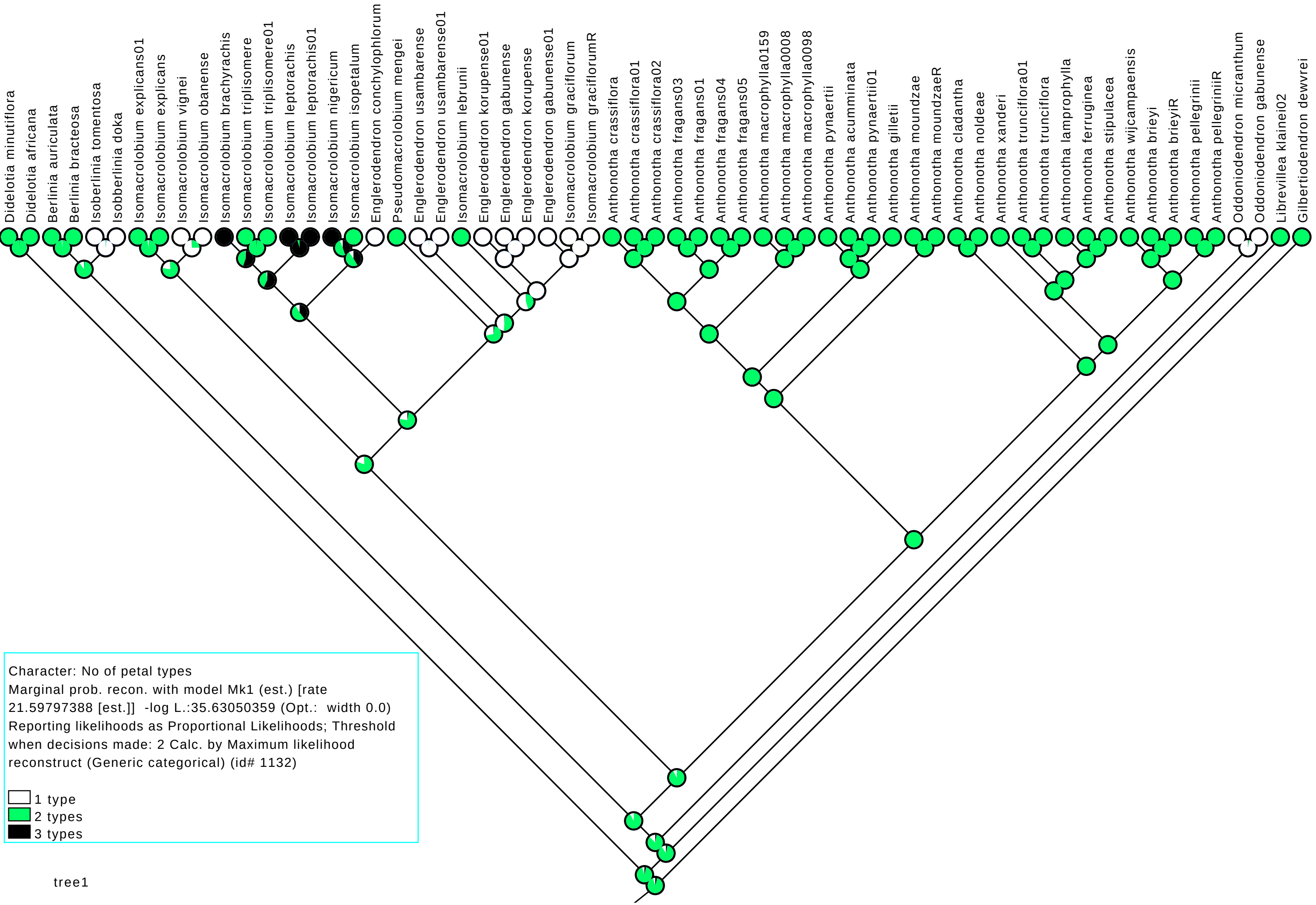
