## Supplementary material for "Phylogenomics within the Anthonotha clade (Detarioideae, Leguminosae) reveals a high diversity in floral trait shifts and a general trend towards organ number reduction": Fig. S7

### Slide 1
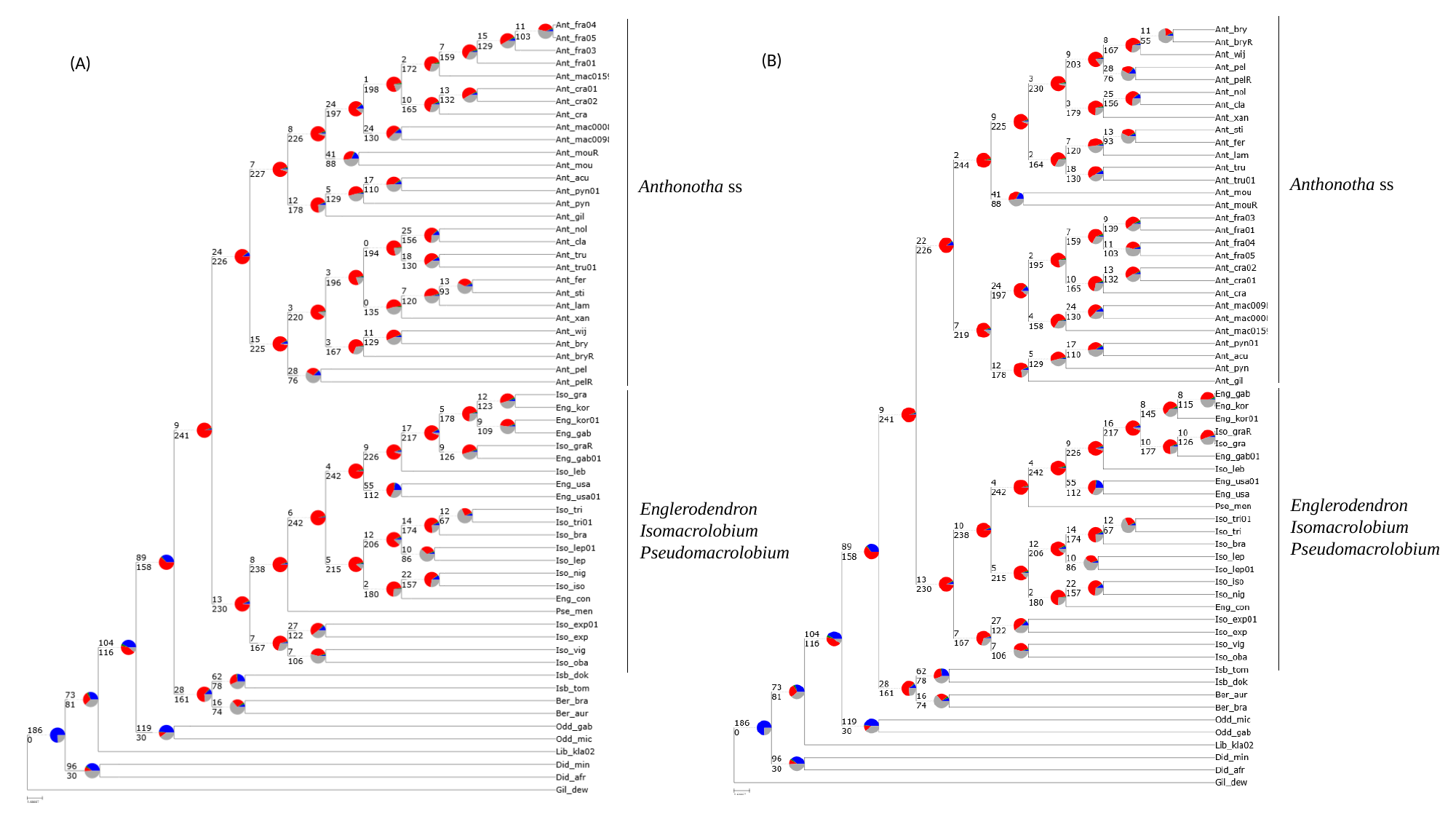

(B)
(A)
Anthonotha ss
Anthonotha ss
Englerodendron
Isomacrolobium
Pseudomacrolobium
Englerodendron
Isomacrolobium
Pseudomacrolobium
