## Supplementary material for "Phylogenomics within the Anthonotha clade (Detarioideae, Leguminosae) reveals a high diversity in floral trait shifts and a general trend towards organ number reduction": Fig. S10

### Slide 1
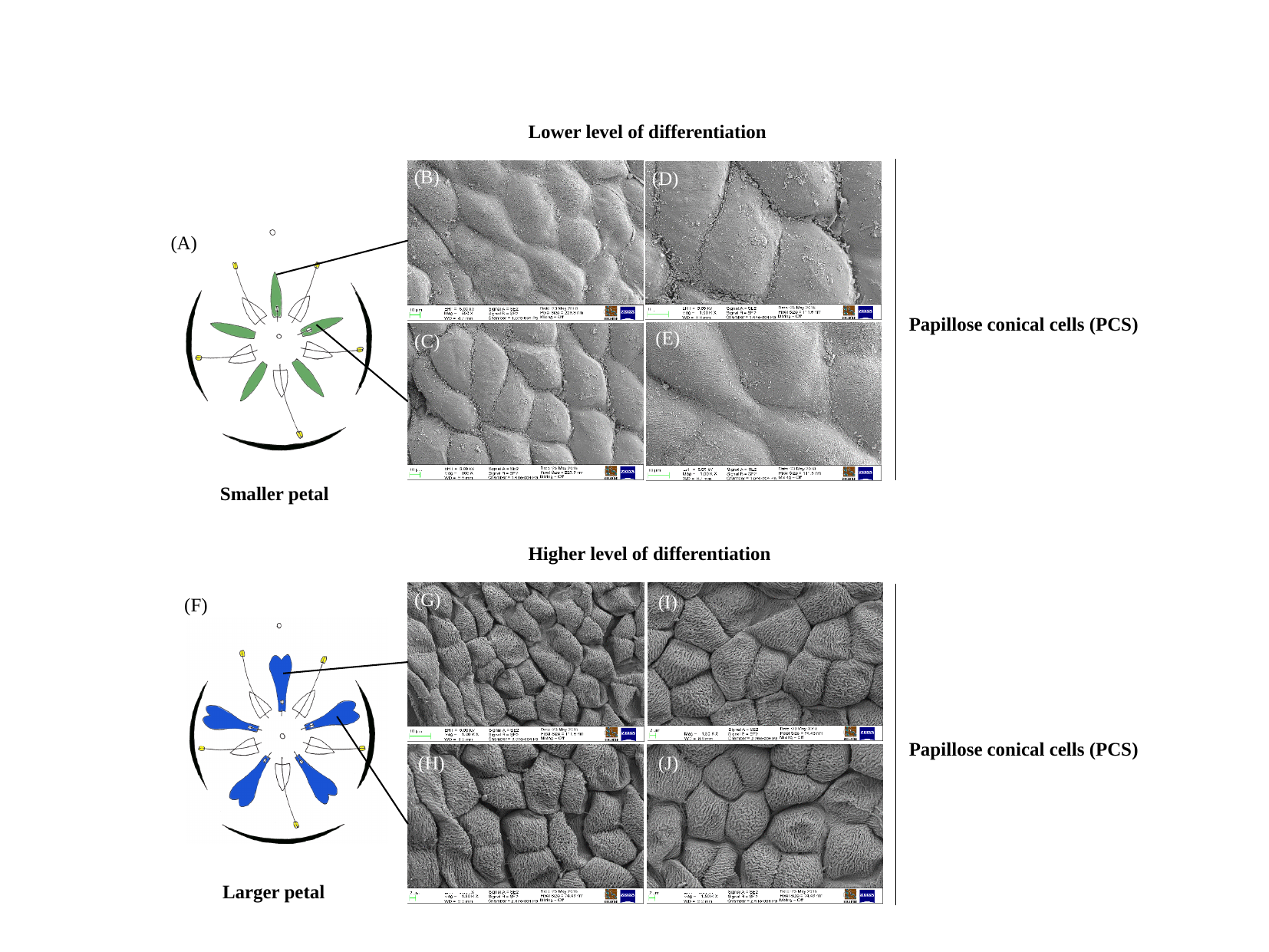

Lower level of differentiation
(B)
(D)
(A)
Papillose conical cells (PCS)
(E)
(C)
Smaller petal
Higher level of differentiation
(G)
(I)
(F)
Papillose conical cells (PCS)
(H)
(J)
Larger petal
