## Supplementary material for "Phylogenomics within the Anthonotha clade (Detarioideae, Leguminosae) reveals a high diversity in floral trait shifts and a general trend towards organ number reduction": Table S1

|  | ***Anthonotha fragans*** | ***Afzelia bella*** | ***Prioria balsamifera*** | ***Copaifera officinalis*** |
| --- | --- | --- | --- | --- |
| **Total lenght of reads (bp)** | 25,259,219 | 22,618,356 | 25,161,399 | - |
| **Total assembled bases (bp)** | 65,929,801 | 45,370,894 | 59,336,292 | 48,588,793 |
| **Total number of contigs** | 86,488 | 60,954 | 75,481 | 52,752 |
| **Average of assembled contig** | 762.3 | 744.35 | 786.11 | 921.08 |
| **Transcripts > 500 bp** | 38,781 | 30,312 | 33,047 | 29,149 |
| **Transcripts > 1000 bp** | 21,216 | 14,785 | 18,829 | 16,946 |
| **Longest transcript (bp)** | 11,522 | 19,361 | 18,844 | 13,545 |
| **Median contig length** | 436 | 497 | 424 | 580.5 |
| **N50 length (bp)** | 1,269 | 1,100 | 1,361 | 1,467 |
| **% GC content** | 42.36 | 41.62 | 41.73 | 41.68 |
| **BUSCO v 2.0.1 embryophyta_odb9** | C:75.3% [S:48.8%, D:26.5%]  F:8.5%, M:16.2%, n:1440 | C:58.9% [S:43.6%, D:15.3%]  F:13.9%, M:27.2%, n:1440 | C:78.2% [S:54.8%, D:23.4%]  F:6.7%, M:15.1%, n:1440 | C:65.6% [S:49.8%, D:15.8%]  F:10.5%, M:23.9%, n:1440 |
| **No. of ORFs identified (predicted)** | 61,138 | 43,453 | 49,878 | 41,788 |
| ***No. of ORFs with BlastP hits** | 46,510 | 35,063 | 37,451 | 32,129 |
