## Supplementary material for "Phylogenomics within the Anthonotha clade (Detarioideae, Leguminosae) reveals a high diversity in floral trait shifts and a general trend towards organ number reduction": Table S2

| **Species** | **Location** | **Voucher specimen (Herbarium)** |
| --- | --- | --- |
| *Gilbertiodendron dewevrei* (De Wild.) J. Léonard |  | *F.J. Breteler 15433* (WAG) |
| *Didelotia africana* Baill. |  | *F.J. Breteler 14374* (WAG) |
| *Didelotia minutiflora* (A. Chev.) J. Léonard |  | *J.J. Wieringa 2424* (WAG) |
| *Librevillea klainei* (Pierre ex Harms) Hoyle |  | *Nkondi 638* (K) |
| *Oddoniodendron micranthum* Baker f. |  | F.J. Breteler 14181 (WAG) |
| *Oddoniodendron gambanum* Ngok & Breteler | GABON | *G. McPherson 16251* (BR) |
| *Isoberlinia tomentosa* (Harms) Craib & Stapf | BENIN, zone cynegetique de la Pendjari, Dassari | Barbier 358 (BR) |
| *Isoberlinia doka* Craib & Stapf | GHANA, Mole national park near headquarters | C.C.H Jongkind 1995 (BR) |
| *Berlinia auriculata* Benth. | CONGO (Brazaville), Lekoumu Prefecture | Bongou, Y. 104 (BR) |
| *Berlinia bracteosa* Benth. | CAMEROON, South, west province, Korup National Park | X.M van der Burgt 748 (BR) |
| *Anthonotha gilletii* (De Wild.) J. Léonard | CONGO K., Kinshasa | Pauwels, L. 5455 (WAG) |
| *Anthonotha pynaertii* (De Wild.) Exell & Hillc. | CONGO K., Bandundu | Devred, R. 2008 (WAG) |
|  | CONGO K., Haut-Ogooué | Wieringa, J.J. 6378 (WAG) |
| *Anthonotha acuminata* (De Wild.) Exell & Hillc. | CAMEROON, South Province | Letouzey, R. 10057 (WAG) |
| *Anthonotha macrophylla* P. Beauv. | CAMEROON, South Province | Andel, T.R. van 3407 (WAG) |
|  | GUINÉE EQUATORIALE, Monte Alen | Ngomo 944 BRLU |
|  | LIBERIA, Grand Cap Mount | Jongkind, C.C.H. 11686 (WAG) |
| *Anthonotha crassifolia* (Baill.) J. Léonard | GABON, Ogooué-Ivindo | McPherson, G.D. 16099 (WAG) |
|  | SIERRA LEONE, Northern Province | Morton, J.K. 2847 (WAG) |
|  | BENIN, Ouémé | Adjakidjè, V. 4702 (WAG) |
| *Anthonotha fragrans* (Baker f.) Exell & Hillc. | CAMEROON, South-West | Burgt, X.M. van der 743 (WAG) |
|  | GABON, Ogooué-Lolo | Wieringa, J.J. 6524 (WAG) |
|  | CÔTE D'IVOIRE, Agboville | Wieringa, J.J. 4291 (WAG) |
|  | LIBERIA, Nimba | Nimba Botanic Team 176 (WAG) |
| *Anthonotha xanderi* Brettler sp. nov | CAMEROON, South-West Province | Burgt, X.M. van der 729 (WAG) |
| *Anthonotha cladantha* (Harms) J. Léonard | CAMEROON, East Province | Letouzey, R. 5621 (WAG) |
| *Anthonotha noldeae* (Rossberg) Exell & Hillc. | CAMEROON, North-West Province | Letouzey, R. 13148 (WAG) |
| *Anthonotha lamprophylla* (Harms) J. Léonard | CAMEROON, South Province | Burgt, X.M. van der 154 (WAG) |
| *Anthonotha ferruginea* (Harms) J. Léonard | GABON, Ogooué-Maritime | Mouandza Mbembo, J.-.C. 127 (WAG) |
| *Anthonotha stipulacea* J. Léonard | GABON, Estuaire | Maas, P.J.M. 10439 (WAG) |
| *Anthonotha trunciflora* J. Léonard | GABON | J.J. Wieringa 1225 (BR) |
|  | GABON, Ogooué-Maritime | Burgt, X.M. van der 77 (WAG) |
| *Anthonotha pellegrinii* Aubrév. | GABON, Estuaire | Floret, J.J. 1395 ((WAG)) |
| *Anthonotha wijcampaensis* Breteler sp. nov | CAMEROON, South Province | Wilde, J.J.F.E. de 8272 (WAG) |
| *Anthonotha brieyi* (De Wild.) J. Léonard | GABON, Ogooué-Lolo | Wieringa, J.J. 6082 (WAG) |
|  | GABON, Ogooué-Lolo *(Replicate)* | Wieringa, J.J. 6082 (WAG) |
| *Anthonotha mouandzae* Breteler | GABON, Ogooué-Maritime | Harris, D.J. 8623 (WAG) |
|  | GABON, Ogooué-Maritime *(Replicate)* | Harris, D.J. 8623 (WAG) |
| *Isomacrolobium obanense* (Baker f.) Aubrév. & Pellegr. | LIBERIA | C.C.H Jongkind, 9748 (BR) |
| *Isomacrolobium vignei* (Hoyle) Audrév. & Pellegr. | LIBERIA, Grand Gedeh, north of Sapo National Park | C.C.H Jongkind 11964 (BR) |
| *Isomacrolobium explicans* (Baill.) Breteler | GUINEE, route Kindia_Telimele, km 20 | Lisowski 51534 (BR) |
|  | *M. Cheek 16972* (K) |  |
| *Isomacrolobium nigericum* (Baker f.) Aubrév. & Pellegr. | DEMOCRATIC REPUBLIC OF CONGO, Kinganga | R. Devred 1252 (BR) |
| *Isomacrolobium isopetalum* (Harms) Audrév. & Pellegr. | CAMEROON, SW of Lelen near Ngoila. | R. Letouzey 11704 (BR) |
| *Isomacrolobium triplisomere* (Pelleg.) Breteler | GABON, Ngounié, 25 km on the road Ikobey to Bakongue | J.J. Wieringa 4439 (BR) |
| *Isomacrolobium brachyrhachis* Breteler | GABON | Louis A.M., Breteler F.J. & De Bruijn J. 729 (BR) |
| *Isomacrolobium leptorachis* (Harms) Aubrév. & Pellegr. | CAMEROON, South Province E slope ofElephant mountain(N’Kol Ebunde | J.J. Wieringa 2140 (BR) |
|  | Rickson s.n. (MT) |  |
| *Isomacrolobium lebrunii* (J. Léonard) Aubrév. & Pellegr. | DEMOCRATIC REPUBLIC OF THE CONGO | L. Lebrun 6497 (HOLOTYPE) (BR) |
| *Isomacrolobium graciflorum* (Harms) Aubrév. & Pellegr. | CONGO, Loukanga II, 25 km de Brazzaville vers | J. Lejoly 86/203 (BR) |
|  | CONGO, Loukanga II, 25 km de Brazzaville vers *(Replicate)* | J. Lejoly 86/203 (BR) |
| *Englerodendron conchyliophorum* Breteler | GABON | Van Valkenburg J.L.C.H 2793A (BR) |
| *Englerodendron usambarense* Harms | P. Herendeen 17-XII-97-8 (USA) |  |
|  | TANZANIA, Tanganika, Kwamkor, Tanga Dist. | S.R. Semsei 2942 (BR) |
| *Englerodendron gabunense* (J. Léonard) Breteler |  | C. Wilks 427 (K) |
|  | GABON, Ogooue-Lolo, c 50 km N of Lastoursville | J.J. Wieringa 6213 (BR) |
| *Englerodendron korupense* Burgt | CAMEROON, South-West Province, Korup National Park. NW plot near P transect, subplot 33C | X. van der Burgt 747 (K) |
| *Pseudomacrolobium mengei* Hauman |  | J. Louis 3666 (K) |
