## Supplementary material for "Phylogenomics within the Anthonotha clade (Detarioideae, Leguminosae) reveals a high diversity in floral trait shifts and a general trend towards organ number reduction": Table S3

| **Species** | **Flower symmetry** | **No. petals** | **No. petal types** | **Petal merism** | **No. of stamens** | **No. of staminoids** | **Stamen merism** |
| --- | --- | --- | --- | --- | --- | --- | --- |
| *Anthonotha acuminata* | 0 | 1 | 1 | 1 | 1 | 1 | 1 |
| *Anthonotha brieyi* | 0 | 2 | 1 | 0 | 1 | 2 | 0 |
| *Anthonotha cladantha* | 0 | 1 | 1 | 1 | 1 | 1 | 1 |
| *Anthonotha crassifolia* | 0 | 1 | 1 | 1 | 1 | 1 | 1 |
| *Anthonotha ferruginea* | 0 | 1 | 1 | 1 | 1 | 1 | 1 |
| *Anthonotha fragrans* | 0 | 1 | 1 | 1 | 1 | 1 | 1 |
| *Anthonotha gilletii* | 0 | 1 | 1 | 1 | 1 | 1 | 1 |
| *Anthonotha lamprophylla* | 0 | 1 | 1 | 1 | 1 | 1 | 1 |
| *Anthonotha macrophylla* | 0 | 1 | 1 | 1 | 1 | 1 | 1 |
| *Anthonotha moundzae* | 0 | 2 | 1 | 0 | 1 | 1 | 1 |
| *Anthonotha noldeae* | 0 | 2 | 1 | 1 | 2 | 2 | 0 |
| *Anthonotha pellegrinii* | 0 | 2 | 1 | 0 | 1 | 2 | 1 |
| *Anthonotha pynaertii* | 0 | 1 | 1 | 1 | 1 | 1 | 1 |
| *Anthonotha stipulacea* | 0 | 1 | 1 | 1 | 1 | 1 | 1 |
| *Anthonotha trunciflora* | 0 | 1 | 1 | 1 | 1 | 1 | 1 |
| *Anthonotha wijmacampensis* | 0 | 2 | 1 | 0 | 1 | 2 | 1 |
| *Anthonotha xanderi* | 0 | 1 | 1 | 1 | 1 | 1 | 1 |
| *Isomacrolobium graciflorum* | 0 | 2 | 0 | 0 | 1 | 2 | 0 |
| *Isomacrolobium explicans* | 0 | 1 | 1 | 1 | 1 | 1 | 1 |
| *Isomacrolobium brachyrachis* | 0 | 1 | 2 | 1 | 1 | 2 | 1 |
| *Isomacrolobium leptorachis* | 0 | 1 | 2 | 1 | 1 | 1 | 1 |
| *Isomacrolobium lebrunii* | 0 | 2 | 1 | 0 | 1 | 1 | 1 |
| *Isomacrolobium obanense* | 0 | 1 | 0 | 1 | 1 | 1 | 1 |
| *Isomacrolobium nigericum* | 0 | 1 | 2 | 1 | 1 | 1 | 1 |
| *Isomacrolobium isopetalum* | 0 | 2 | 1 | 1 | 1 | 2 | 0 |
| *Isomacrolobium triplesomere* | 0 | 1 | 1 | 1 | 1 | 1 | 1 |
| *Isomacrolobium vignei* | 0 | 2 | 0 | 1 | 1 | 2 | 1 |
| *Englerodendron korupense* | 1 | 1 | 0 | 1 | 2 | 2 | 1 |
| *Englerodendron gabunense* | 1 | 1 | 0 | 1 | 2 | 2 | 1 |
| *Englerodendron conchyliophorum* | 1 | 1 | 0 | 1 | 2 | 2 | 1 |
| *Englerodendron usambarense* | 1 | 0 | 0 | 1 | 2 | 1 | 1 |
| *Pseudomacrolobium mengei* | 0 | 1 | 1 | 1 | 0 | 0 | 1 |
| *Berlinia auriculata* | 0 | 1 | 1 | 1 | 0 | 0 | 1 |
| *Berlinia bracteosa* | 0 | 1 | 1 | 1 | 0 | 0 | 1 |
| *Isoberlinia tomentosa* | 1 | 1 | 0 | 1 | 0 | 0 | 1 |
| *Isoberlinia doka* | 1 | 1 | 0 | 1 | 0 | 0 | 1 |
| *Oddoniodendrum gabunense* | 1 | 1 | 0 | 1 | 0 | 0 | 1 |
| *Oddoniodendron micranthum* | 1 | 1 | 0 | 1 | 0 | 0 | 1 |
| *Librevillea klainei* | 1 | 3 | 1 | 1 | 0 | 0 | 1 |
| *Didelotia africana* | 0 | 1 | 1 | 1 | 0 | 2 | 1 |
| *Didelotia minutiflora* | 0 | 1 | 1 | 1 | 0 | 2 | 1 |
| *Gilbertiodendron dewevrei* | 0 | 1 | 1 | 1 | 1 | 1 | 1 |
| Flower symmetry, zygomorphic (0), actinomorphic (1); No. petals, 6 petals (0), five petals (1), 1-4 petals (2), absent (3); No. petal types, one type (0), two types (1), three types (2); intraspecific petal merism, variable (0), invariable (1); fertile stamens, >6 (0), 3 (1), 6-5 (2); staminoids, absent (0), six (1) between 1-5 6 (2); intraspecific stamen merism, variable (0), invariable (1) | | | | | | | |
