## Supplementary material for "Phylogenomics within the Anthonotha clade (Detarioideae, Leguminosae) reveals a high diversity in floral trait shifts and a general trend towards organ number reduction": Table S4

| **Ancestral in *Anthonotha* clade** | **Voucher specimen** | **Classification of petal types based on location on the dorsoventral axis within the flower** | | |
| --- | --- | --- | --- | --- |
|  |  | **Adaxial** | **Lateral** | **Abaxial** |
| *Anthonotha brieyii* | G. Le Testu 1737 | PCS^t^ | PCS^t^ | PCS^t^ |
| *Anthonotha pynaertii* | Gilbert 10297 | PKR^t^ | PKR^t^ | PKR^t^ |
| *Anthonotha acuminata* | De Wilde 7623 | PKR^t^ | PKR^t^ | PKR^t^ |
| *Anthonotha noldeae* | Michelson 1060, Kahurangnga 2635 | PKR^t^ | PKR^t^ | PKR^t^ |
| *Anthonotha lamprohylla* | Thomas 6908 | PKR | PKR | PKR |
| *Anthonotha stipulacea* | Dauby 1134 | PKR | PKR | PKR |
| *Anthonotha macrophylla* | Louis 13570, Pawells 6873 | PKR | PKR | PKR |
| *Anthonotha fragans* | M. Leal 178 | PCS | PCS | PCS |
| *Anthonotha gilletii* | Devred 2159 | PKR | PKR | PKR |
| *Anthonotha crassifolia* | D.N. Thocea 4698 | PKR | PKR | PKR |
| *Isomacrolobium isopetalum* | Letovzcfy 11704 | PCS | PCS | PCS |
| **Reduction to one petal type** | | | | |
| *Englerodendron conchylophlorum* | Carvahlo 6046 | PCS | PCS | PCS |
| *Englerodendron korupense* | Van Der Burgt 741 | PCS | PCS | PCS |
| *Englerodendron usambarense* | Semsei 2942 | PCS^t^ | PCS^t^ | PCS^t^ |
| *Englerodendron gabunense* | Bretteler 11081 | PKR | PKR | PKR |
| *Isomacrolobium obanense* | Jongkind 9748 | PKR | PKR | PKR |
| *Isomacrolobium graciflorum* | Lejoly 86/203 | PKR | PKR | PKR |
| *Isomacrolobium vignei* | Dewilde 3663 | PCS | PCS | PCS |
| **Same petal number type but alternative arrangement** | | | | |
| *Pseudomacrolobium mengei* | Leonard 721 | PKR | PKR | PKR |
| *Isomacrolobium explicans* | S. Lisowaski 54235 | PKR | PKR | PKR |
| **Increase to three petal types** | | | | |
| *Isomacrolobium leptorhachis* | J.J. Boss 3290 | PCS | PCS | PCS |
