## Supplementary material for "Phylogenomics within the Anthonotha clade (Detarioideae, Leguminosae) reveals a high diversity in floral trait shifts and a general trend towards organ number reduction": Table S5

| **Species** | **No. of reads** | **No. of reads mapped** | **% reads mapped** | **% of bait captured** | **Assembled contigs (gSPADes)** |
| --- | --- | --- | --- | --- | --- |
| *Anthonotha acuminata* | 472,140 | 200,674 | 42.51 | 88.6 | 9,365 |
| *Anthonotha brieyi* | 292,492 | 130,295 | 44.55 | 86.4 | 6,173 |
| *Anthonotha brieyi_R* | 515,657 | 77,516 | 15.03 | 73.6 | 2,548 |
| *Anthonotha cladantha* | 1,034,215 | 394,958 | 38.19 | 85.6 | 6,379 |
| *Anthonotha crassifolia* | 1,033,775 | 439,332 | 42.51 | 85.71 | 6,156 |
| *Anthonotha crassifolia01* | 707,572 | 308,542 | 43.61 | 86.8 | 7,501 |
| *Anthonotha crassifolia02* | 945,733 | 417,438 | 44.14 | 88.2 | 9,690 |
| *Anthonotha ferruginea* | 723,687 | 323,339 | 44.68 | 89.29 | 13,396 |
| *Anthonotha fragrans01* | 986,512 | 393,593 | 39.91 | 85.7 | 7,564 |
| *Anthonotha fragrans03* | 893,807 | 366,338 | 40.99 | 87.74 | 10,526 |
| *Anthonotha fragrans04* | 836,066 | 366,259 | 43.81 | 88.96 | 13,829 |
| *Anthonotha fragrans05* | 359,558 | 149,589 | 41.59 | 87.59 | 8,691 |
| *Anthonotha gilletii* | 703,889 | 292,697 | 41.58 | 83.55 | 3,647 |
| *Anthonotha lamprophylla* | 758,280 | 331,647 | 43.74 | 88.25 | 12,629 |
| *Anthonotha macrophylla* _*BoD0098* | 947,697 | 419,102 | 44.22 | 86.26 | 7,721 |
| *Anthonotha macrophylla* _*BoD0008* | 655,930 | 331,627 | 50.56 | 83.25 | 2,522 |
| *Anthonotha macrophylla* *_BoD0159* | 723,385 | 395,954 | 54.74 | 88.19 | 7,752 |
| *Anthonotha mouandzae* | 247,421 | 113,577 | 45.9 | 85.25 | 4,204 |
| *Anthonotha mouandzae_R* | 1,828,659 | 308,728 | 16.88 | 82 | 3,451 |
| *Anthonotha noldeae* | 1,111,414 | 445,343 | 40.07 | 85.13 | 6,036 |
| *Anthonotha pellegrinii* | 192,622 | 73,337 | 38.07 | 77.13 | 1,638 |
| *Anthonotha pellegrinii_R* | 1,580,521 | 231,721 | 14.66 | 81.09 | 5,493 |
| *Anthonotha pynaertii* | 1,004,677 | 400,433 | 39.86 | 86.11 | 6,517 |
| *Anthonotha pynaertii01* | 557,332 | 212,741 | 38.17 | 88.21 | 13,704 |
| *Anthonotha stipulacea* | 1,036,364 | 441,012 | 42.55 | 90.5 | 22,738 |
| *Anthonotha trunciflora* | 1,161,113 | 570,646 | 49.15 | 87.67 | 9,338 |
| *Anthonotha trunciflora01* | 2,115,844 | 979,911 | 46.31 | 89.36 | 19,406 |
| *Anthonotha wijcampaensis* | 2,594,152 | 967,860 | 37.31 | 86.6 | 13,521 |
| *Anthonotha xanderi* | 1,156,424 | 488,566 | 42.25 | 89.67 | 18,172 |
| *Isomacrolobium graciflorum* | 405,613 | 172,334 | 42.49 | 86.04 | 6,725 |
| *Isomacrolobium graciflorum_R* | 2,260,559 | 464,941 | 20.57 | 83.3 | 4,635 |
| *Isomacrolobium explicans* | 495,003 | 200,153 | 40.43 | 85.23 | 4,813 |
| *Isomacrolobium explicans01* | 444,055 | 100,284 | 22.58 | 81.52 | 5,164 |
| *Isomacrolobium brachyrachis* | 1,746,137 | 642,911 | 36.82 | 86.93 | 11,741 |
| *Isomacrolobium leptorachis* | 463,830 | 204,263 | 44.04 | 88.02 | 10,551 |
| *Isomacrolobium leptorachis01* | 621,536 | 126,201 | 20.31 | 81.54 | 5,269 |
| *Isomacrolobium lebrunii* | 1,890,244 | 181,893 | 9.62 | 76.86 | 2,609 |
| *Isomacrolobium obanense* | 410,185 | 186,490 | 45.46 | 88.41 | 8,319 |
| *Isomacrolobium nigericum* | 1,313,251 | 587,793 | 44.76 | 86.07 | 6,784 |
| *Isomacrolobium isopetalum* | 960,648 | 411,799 | 42.87 | 85.17 | 5,142 |
| *Isomacrolobium triplesomere* | 633,826 | 288,624 | 45.54 | 88.8 | 11,159 |
| *Isomacrolobium triplesomere01* | 680,650 | 164,350 | 24.15 | 81.16 | 4,137 |
| *Isomacrolobium vignei* | 475,978 | 226,649 | 47.62 | 88.63 | 10,880 |
| *Englerodendron korupense* | 610,918 | 292,419 | 47.87 | 88.5 | 10,285 |
| *Englerodendron korupense01* | 1,213,111 | 276,534 | 22.81 | 84.89 | 7,468 |
| *Englerodendron gabunense* | 374,708 | 175,935 | 46.95 | 87.86 | 6,469 |
| *Englerodendron gabunense01* | 2,510,336 | 445,192 | 17.73 | 82.08 | 3,664 |
| *Englerodendron conchyliophorum* | 565,591 | 258,974 | 45.79 | 88.45 | 11,762 |
| *Englerodendron usambarense* | 1,653,147 | 723,777 | 43.78 | 86.36 | 7,541 |
| *Englerodendron usambarense01* | 935,334 | 187,886 | 20.09 | 81.83 | 6,379 |
| **Outgroup taxa** |  |  |  |  |  |
| *Pseudomacrolobium mengei* | 1,550,479 | 498,404 | 32.15 | 79.96 | 2,294 |
| *Berlinia auriculata* | 421,586 | 200,092 | 47.46 | 88.23 | 9,478 |
| *Berlinia bracteosa* | 556,139 | 111,786 | 20.11 | 79.18 | 3,261 |
| *Isoberlinia tomentosa* | 276,398 | 127,964 | 46.3 | 85.61 | 4,652 |
| *Isoberlinia doka* | 704,085 | 186,260 | 26.45 | 83.46 | 6,417 |
| *Oddoniodendrum gabunense* | 298,224 | 154,010 | 51.64 | 83.87 | 3,600 |
| *Oddoniodendron micranthum* | 911,753 | 261,536 | 28.68 | 85.19 | 10,012 |
| *Librevillea klainei* | 347,552 | 29,735 | 8.56 | 72.57 | 6,112 |
| *Didelodia africana* | 1,104,456 | 73,732 | 6.68 | 78.88 | 9,333 |
| *Didelodia minutiflora* | 1,320,101 | 97,069 | 7.35 | 81.05 | 13,817 |
| *Gilbertiodendron dewevrei* | 778,876 | 55,698 | 7.15 | 80.25 | 19,300 |
