## Supplementary material for "Phylogenomics within the Anthonotha clade (Detarioideae, Leguminosae) reveals a high diversity in floral trait shifts and a general trend towards organ number reduction": Table S6

| **Species** | **No. orthologues** | **Size (bp)** | **% of orthologues** | **% of size (bp)** |
| --- | --- | --- | --- | --- |
| Ant_acu | 694 | 192,380 | 0.752711496746 | 0.803813916953 |
| Ant_bry | 711 | 189,611 | 0.77114967462 | 0.792244311297 |
| Ant_bryR | 537 | 122,073 | 0.582429501085 | 0.510052896789 |
| Ant_cla | 738 | 184,977 | 0.800433839479 | 0.772882248239 |
| Ant_cra | 740 | 187,995 | 0.802603036876 | 0.785492240969 |
| Ant_cra01 | 743 | 193,939 | 0.805856832972 | 0.810327826385 |
| Ant_cra02 | 713 | 195,412 | 0.773318872017 | 0.816482405341 |
| Ant_fer | 705 | 195,399 | 0.76464208243 | 0.816428087944 |
| Ant_fra01 | 720 | 184,438 | 0.780911062907 | 0.770630165376 |
| Ant_fra03 | 730 | 194,968 | 0.791757049892 | 0.814627257306 |
| Ant_fra04 | 697 | 192,516 | 0.755965292842 | 0.804382160495 |
| Ant_fra05 | 687 | 188,852 | 0.745119305857 | 0.789073010939 |
| Ant_gil | 712 | 175,334 | 0.772234273319 | 0.732591274119 |
| Ant_lam | 719 | 195,044 | 0.779826464208 | 0.814944805168 |
| Ant_mac0008 | 719 | 173,268 | 0.779826464208 | 0.723958986187 |
| Ant_mac0098 | 737 | 191,115 | 0.799349240781 | 0.798528416355 |
| Ant_mac0159 | 715 | 194,038 | 0.775488069414 | 0.810741474258 |
| Ant_mou | 716 | 186,194 | 0.776572668113 | 0.777967192292 |
| Ant_mouR | 670 | 156,629 | 0.726681127983 | 0.654436895719 |
| Ant_nol | 751 | 185,711 | 0.81453362256 | 0.775949092064 |
| Ant_pel | 579 | 137,392 | 0.627982646421 | 0.574059682285 |
| Ant_pelR | 671 | 154,285 | 0.727765726681 | 0.644643051134 |
| Ant_pyn | 716 | 184,503 | 0.776572668113 | 0.770901752363 |
| Ant_pyn01 | 715 | 192,524 | 0.775488069414 | 0.804415586586 |
| Ant_sti | 669 | 187,029 | 0.725596529284 | 0.781456040512 |
| Ant_tru | 733 | 193,381 | 0.795010845987 | 0.807996356556 |
| Ant_tru01 | 731 | 197,653 | 0.79284164859 | 0.825845889009 |
| Ant_wij | 744 | 189,732 | 0.80694143167 | 0.79274988092 |
| Ant_xan | 689 | 192,027 | 0.747288503254 | 0.802338990699 |
| Ber_aur | 712 | 192,948 | 0.772234273319 | 0.806187169395 |
| Ber_bra | 643 | 151,009 | 0.697396963124 | 0.630955066978 |
| Did_afr | 630 | 142,682 | 0.683297180043 | 0.596162684784 |
| Did_min | 640 | 153,049 | 0.694143167028 | 0.639478720115 |
| Eng_con | 708 | 193,849 | 0.767895878525 | 0.809951782864 |
| Eng_gab | 707 | 190,918 | 0.766811279826 | 0.797705298871 |
| Eng_gab01 | 646 | 145,212 | 0.700650759219 | 0.606733685979 |
| Eng_kor | 717 | 192,725 | 0.777657266811 | 0.805255417116 |
| Eng_kor01 | 708 | 177,908 | 0.767895878525 | 0.743346118813 |
| Eng_usa | 751 | 188,731 | 0.81453362256 | 0.788567441316 |
| Eng_usa01 | 704 | 167,890 | 0.763557483731 | 0.70148829669 |
| Gil_dew | 664 | 162,278 | 0.720173535792 | 0.678039894039 |
| Isb_dok | 701 | 175,017 | 0.760303687636 | 0.731266765274 |
| Isb_tom | 698 | 181,307 | 0.75704989154 | 0.757548029114 |
| Iso_bra | 733 | 190,661 | 0.795010845987 | 0.796631485706 |
| Iso_exp | 694 | 178,231 | 0.752711496746 | 0.744695697226 |
| Iso_exp01 | 678 | 166,173 | 0.73535791757 | 0.694314221966 |
| Iso_gra | 745 | 190,095 | 0.808026030369 | 0.794266589787 |
| Iso_graR | 717 | 170,603 | 0.777657266811 | 0.712823919711 |
| Iso_iso | 735 | 183,303 | 0.797180043384 | 0.765887838753 |
| Iso_leb | 428 | 98,340 | 0.46420824295 | 0.410890220362 |
| Iso_lep | 701 | 193,397 | 0.760303687636 | 0.808063208738 |
| Iso_lep01 | 668 | 164,186 | 0.724511930586 | 0.68601201668 |
| Iso_nig | 746 | 188,446 | 0.809110629067 | 0.787376636834 |
| Iso_oba | 707 | 194,306 | 0.766811279826 | 0.811861248297 |
| Iso_tri | 709 | 194,973 | 0.768980477223 | 0.814648148612 |
| Iso_tri01 | 662 | 158,280 | 0.718004338395 | 0.661335205194 |
| Iso_vig | 706 | 193,486 | 0.765726681128 | 0.808435073997 |
| Lib_kla02 | 526 | 120,783 | 0.570498915401 | 0.504662939658 |
| Odd_gab | 690 | 177,987 | 0.748373101952 | 0.743676201459 |
| Odd_mic | 707 | 183,453 | 0.766811279826 | 0.766514577954 |
| Pse_men | 636 | 142,743 | 0.689804772234 | 0.596417558725 |
